# MECP2 Duplication Uncouples Mitochondrial and Purine Metabolism During neuronal maturation

**DOI:** 10.64898/2025.12.24.696399

**Authors:** Gerarda Cappuccio, Guantong Qi, Xuan Qin, Saleh Mahmud Khalil, John Hunyara, Yang Li, Joel Mathews, Armand Soriano, Sivan Osenberg, Senghong Sing, Toni Claire Tacorda, Luke Parkitny, Jennifer Sheppard, George Timpone, Sergio Attanasio, Sara Bitar, Ashley Grace Anderson, Demian Rocha Ifa, Christine Coquery, Bernard Suter, Davut Pehlivan, Hu Chen, Zhandong Liu, Feng Li, Huda Y. Zoghbi, Paymaan Jafar-Nejad, Mirjana Maletic-Savatic

## Abstract

Mitochondria and nucleotide metabolism are critical for cellular and developmental homeostasis, yet their potential interdependence and role in neurodevelopmental disease remain unclear. In *MECP2* Duplication Syndrome (MDS), we identify a conserved correlation between mitochondrial function and purine metabolism that is disrupted across human, organoid, and mouse models. Multiomics integration reveals Complex III as the focal point of mitochondrial collapse, leading to redox stress, DNA damage, and hyperactivation of the *de novo* purine biosynthesis via purinosome assembly. The breakdown of mitochondria–purinosome coupling compromises genome stability, impairs radial glia proliferation, and delays neuronal maturation. By linking a defined genetic dosage imbalance to metabolic network failure, our study positions the mitochondria-purinosome coordination as a fundamental control axis for neurodevelopment and a therapeutic entry point across metabolic and neurodevelopmental disorders. Metabolic control is fundamental to cellular function, influencing energy production, signaling, epigenetic regulation, and tissue homeostasis^1^. Nowhere is this more critical than in the brain, where tightly regulated metabolic networks sustain high energetic demands and support neuronal development, synaptic plasticity, and circuit formation^2^. Disruptions in these networks are increasingly implicated across a spectrum of neurodevelopmental disorders^3–5^, yet their precise metabolic signatures and mechanistic contributions remain poorly understood.

---

Among metabolic regulators, mitochondria occupy a uniquely pivotal position, functioning not only as the primary source of energy but also as integrators of redox balance, biosynthesis, and signaling pathways that govern cell fate decisions^6^. Indeed, mitochondrial dysfunction is increasingly recognized as a convergent feature across diverse neurodevelopmental and neurodegenerative disorders^7–9^.

Despite this central role, how specific genetic mutations reprogram brain metabolism—and whether these changes are causal or secondary—remains unresolved. Genetic disorders offer a powerful lens to address this question because they allow dissection of defined molecular perturbations in human development and physiology. The X-linked transcriptional regulator *MECP2* exemplifies this principle. Altered *MECP2* dosage underlies two severe neurodevelopmental syndromes: Rett syndrome, caused by loss-of-function mutations^10^, and *MECP2* Duplication Syndrome [MDS or X-linked intellectual developmental disorder Lubs type (MRXSL; MIM: 300260)]^11,12^, caused by gene overexpression. Both produce overlapping phenotypes—intellectual disability, epilepsy, motor dysfunction, and early mortality—yet through opposite dosage changes, suggesting that *MECP2* dosage itself destabilizes core cellular homeostasis^13–16^.

*MECP2* is a chromatin regulator that binds methylated and hydroxymethylated CpG dinucleotides, shaping transcriptional and epigenetic landscapes^17^. Through interactions with chromatin-remodeling and histone deacetylase complexes, *MECP2* coordinates gene networks governing neuronal differentiation, synaptic maturation, and activity-dependent plasticity^4–6^. Because *MECP2* encodes two isoforms—*MECP2_e1* (neuronal) and *MECP2_e2* (broadly expressed)^18^—altered dosage likely disrupts both neuronal and systemic metabolism. Indeed, metabolic abnormalities, oxidative stress, impaired ATP production, and impaired mitochondrial morphology, biogenesis, and proteostasis have been observed in Rett syndrome^19–28^, but the metabolic consequences of MECP2 overexpression remain largely unknown.

We hypothesized that *MECP2* overexpression disrupts cellular metabolism in a manner that fundamentally impairs neurodevelopment. To test this, we used an integrated multi-omics strategy spanning patient blood and cerebrospinal fluid (CSF), patient-derived cortical organoids, and a transgenic MDS mouse model in which *MECP2* overexpression was normalized by antisense oligonucleotides (ASOs). Across all cross-species and cross-tissue datasets, we identified convergent disruptions in mitochondrial metabolism and purine biosynthesis, both partially reversible with ASO treatment. In MDS cortical organoids, structural, functional, and metabolic analyses revealed profound mitochondrial deficits centered on the Complex III (CIII) of the electron transport chain (ETC). These deficits were tightly coupled to aberrant *de novo* purine biosynthesis and assembly of purinosomes—dynamic, multi-enzyme condensates that generate nucleotides essential for DNA and RNA synthesis, energy metabolism, and signaling, thereby maintaining genome stability and cellular equilibrium^29,30^. This dysfunctional mitochondria–purinosome axis links mitochondrial collapse to impaired genome maintenance and defective neurogenesis.

Together, these findings establish mitochondrial dysfunction as a primary driver of neurodevelopmental outcomes in MDS—uncovering a unifying mechanism where altered *MECP2* dosage reconfigures metabolism to derail cortical development.

## Cross-Species, Multi-Tissue Multiomics Identifies Conserved Mitochondrial and Nucleotide Pathway Disruptions in MDS

To capture the systemic metabolic consequences of *MECP2* overexpression, we implemented a rigorously controlled, cross-species, multi-model design encompassing human plasma and CSF, patient-derived cortical organoids, and a validated MDS mouse model^31^ treated with ASO targeting *MECP2* transcripts^16^ (**Fig. 1A**, **Extended Data Fig. 1**, **Tables 1–3**). Blood metabolomics from 30 patients with genetically confirmed MDS (all males; 10 sampled twice, 6–8 months apart) were compared to matched parental controls (n=32), minimizing genetic and environmental confounders. Parallel profiling in MDS (n=35) and wild-type (WT) littermate mice robust separation of metabolic profiles (**Fig. 1B**). Across both species, 248 metabolites were consistently altered, including tricarboxylic acid (TCA) cycle intermediates, amino acid derivatives, glutathione metabolism, and purine/pyrimidine nucleotides—implicating broad mitochondrial and redox dysfunction (**Fig. 1D, Extended Data Fig. 2)**. Targeted metabolomics validated these increases (**Extended Data Fig. 1)** and age-stratified analyses together with cross-species validation confirmed that the observed metabolic alterations were not age-dependent (**Extended Data Fig. 2, Table 4)**.

**Figure 1.**
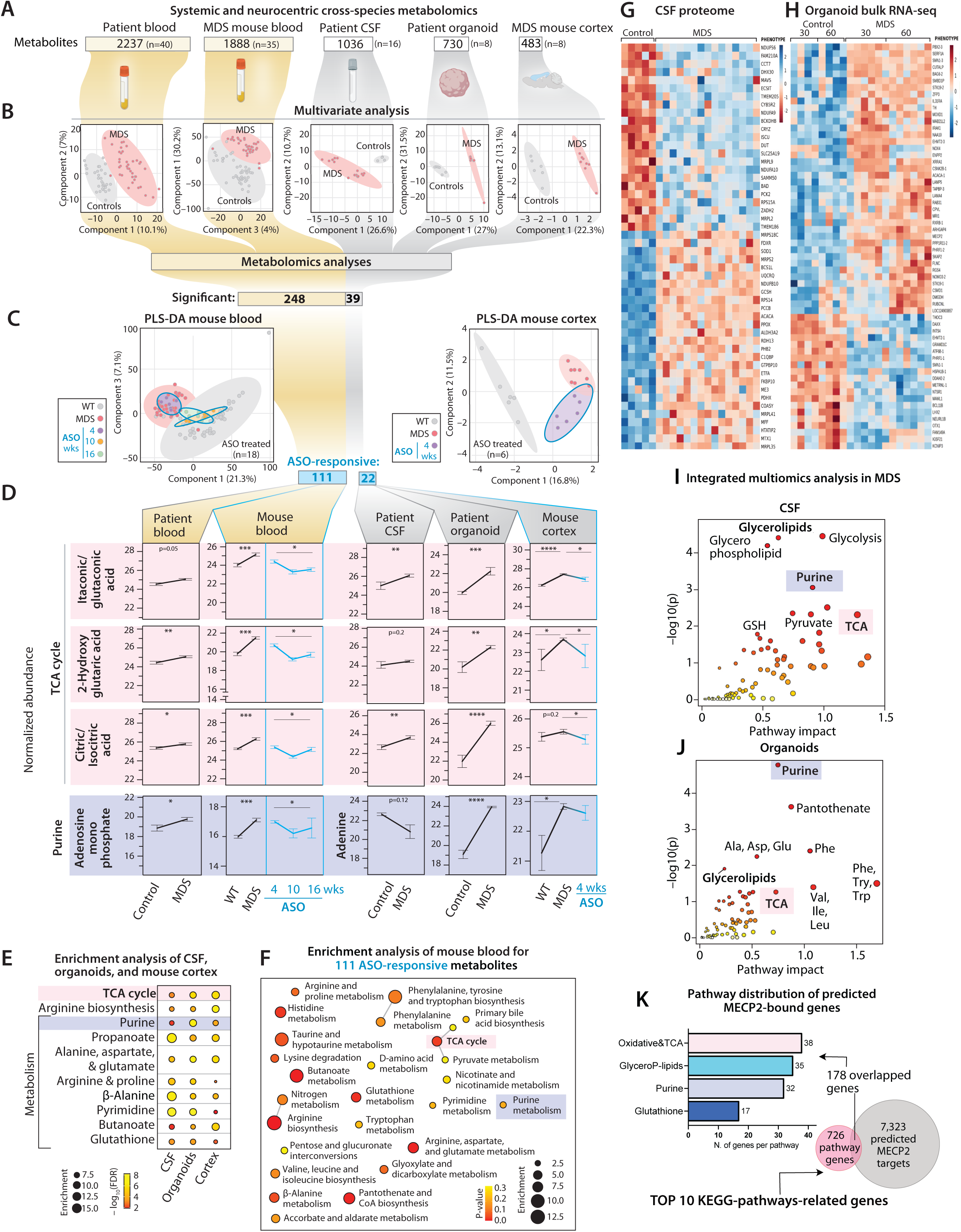
Cross-species, multi-tissue multiomics identifies conserved mitochondrial and nucleotide pathway disruptions in MDS. **A.** Untargeted metabolomics was performed on blood from patients with *MECP2* Duplication Syndrome (MDS) (n=40 from 30 patients; 10 patients sampled twice, 6-8 months apart; n=32 controls (healthy parents) and MDS mouse model (n=35; n=48 wild-type littermates (WT)) (“systemic dataset”) as well as from neural tissues—cerebrospinal fluid (CSF) from MDS patients (n=16; 8 patients sampled twice, 6-8 months apart; n=5 controls (children who required spinal tap as part of the clinical assessment but who did not have any genetic or neurodevelopmental issues), 1 sample run twice), cortical organoids from the subset of MDS patients (n=4 MDS lines, n=3 control lines, two independent batches per line), and cortical tissue from the MDS mice (n=8; n=6 WT controls) (“neurocentric dataset”). All samples were processed on the same LC-MS platform, using the same sample preparation, data collection, and analysis approach. **B.** Partial Least Squares–Discriminant Analysis (PLS-DA) from each dataset shows distinct clustering between MDS and control samples, with each dot representing an individual sample. 248 systemic metabolites and 39 neurocentric metabolites significantly differed between MDS and controls (t-test, p<0.05). **C.** Untargeted metabolomics was performed on blood and cortical tissues from male MDS mice treated with 350 µg of humanized antisense oligonucleotides (ASO) selectively targeting the human *MECP2* copy and analyzed at different timepoints post-treatment (4, 10, and 16 weeks for blood (n=18) and 4 weeks for cortices (n=6)). All samples were processed on the same LC-MS platform in batch runs. PLS-DA shows distinct clustering of the WT, MDS, and ASO-treated MDS across all timepoints. Each dot represents a sample. In both tissues, ASO led to metabolome shift toward controls, with the 10- and 16-week timepoints for blood metabolome overlapping with the WT. 111/248 blood metabolites and 22/39 cortex metabolites responded to ASO toward normalization (one-way ANOVA with post hoc analysis, p<0.05). **D.** Selected metabolites illustrate significant differences across species and tissues, and rescue by ASO. Box-plots are mean±SEM of log2 transformed intensity (*p<0.05, **p<0.01, ***p<0.001, ****p<0.001, based on the meta-analysis combination approach using the **Stouffer method**. For ASO rescue pattern Hunter (2-4-3-2-2 or 4-2-3-4-4 for blood and 1-2-1/2-1-2 for cortex with a correlation >0.25, *p<0.05) was applied. Lines connecting box-plots are for easier visualization of the differences. **E.** Pathway enrichment analysis of the neurocentric dataset (human CSF and organoids, mouse cortex) shows significant enrichment of the TCA cycle, arginine biosynthesis, and purine metabolism as the top three deregulated KEGG pathways (hypergeometric test p<0.05). **F.** Pathway enrichment analysis of 111 ASO-responsive systemic metabolites shows TCA cycle and nucleotide metabolism (hypergeometric test, p<0.05). **G.** CSF proteomics heat map shows top 50 significantly deregulated proteins in MDS. **H.** Heat map of differentially expressed genes (DEGs) derived from bulk RNA sequencing (RNAseq) of 30- and 60-day-old MDS organoids. **I.** Integration of the significantly deregulated CSF metabolites and proteins in MDS highlights shared enrichment of specific pathways including purine metabolism and the TCA cycle (p<0.05, impact > 0.6). **J.** Integration of the significantly deregulated organoid metabolites and transcripts in MDS highlights shared enrichment of specific pathways including purine metabolism and the TCA cycle (p<0.05, impact > 0.6). **K.** Bioinformatic analysis predicted MECP2 binding to the promoter regions of genes associated with deregulated metabolic pathways. 7,323 genes were identified as putative MECP2 targets. Among 726 genes from the CSF and organoid integrative analysis in top 10 KEGG pathways, 178 genes overlapped between predicted MECP2 targets and pathway-associated genes. TCA cycle and oxidative phosphorylation, glycerophospholipid metabolism, purine and glutathione metabolism represent the most enriched categories. Please see **Extended Data** Figures 1-4 and **Tables 1-21** for details.

To determine causality, we treated male MDS mice with 350 µg of humanized ASO selectively targeting the human *MECP2* copy and analyzed plasma at 4-, 10-, and 16-weeks post-treatment. ASO effectively reduced *MECP2* expression toward WT levels (**Extended Data Fig. 1**) and progressively normalized 111 of 248 conserved MDS-associated metabolites in plasma (**Fig. 1C**, **Extended Data Fig. 2, Tables 5-9**). Among these were isocitrate, citrate, 2-hydroxyglutarate (2HG), carnitine derivatives, and purine metabolites (adenine, uracil, uric acid) reflecting restoration of mitochondrial-nucleotide coupling (**Fig. 1D, Extended Data Fig. 2**).

Because MECP2_e1 predominates in neurons whereas MECP2_e2^18^ is expressed across peripheral tissues, we extended our analyses to neurocentric compartments. CSF from a subset of MDS patients (n=8, each sampled twice 6–8 months apart), patient-derived cortical organoids (n=4 lines, 2 batches per line analyzed)^32,33^, and cortical tissues from MDS mice (n=8) revealed convergent alterations in TCA cycle, arginine and purine metabolism (**Fig. 1B, D**)—mirroring plasma findings (**Fig. 1D-F; Extended Data Fig. 3**). Twenty-two of 39 MDS-associated neurocentric metabolites normalized with ASO (**Fig. 1C, D**), including TCA cycle intermediates (isocitrate, citrate), itaconate/glutaconate, 2HG, carnitine derivatives, and purine metabolites (adenine, uracil, uric acid) (**Fig. 1E-F**; **Extended Data Figs. 3**, **Tables 11-15**), confirming reversibility.

To contextualize these metabolic defects within broader molecular networks, we integrated CSF proteomics and cortical organoid transcriptomics. CSF proteomics revealed downregulation of ATP-related pathways and TCA cycle enzymes (**Fig. 1G**), while organoid transcriptomes showed suppressed mitochondrial and glutathione detoxification pathways and induction of oxidative stress responses (**Fig. 1H, Extended Data Fig. 4**). Joint pathway analysis of gene/protein expression with metabolite abundance converged the TCA cycle, purine biosynthesis, and glycerophospholipid metabolism as the core dysregulated modules (**Fig. 1I, J**).

To further examine causality, we asked whether MECP2 directly regulates genes of the most enriched metabolic pathways. Among 726 genes in top ten pathways identified by gene/protein–metabolome integrative analysis (**Extended Data Tables 16–19),** 178 contained predicted *MECP2*-binding domains. Pathways with the highest motif density included the TCA cycle and oxidative phosphorylation (OXPHOS), glycerophospholipid, purine and glutathione metabolism (**Fig. 1K**). Thus, MECP2 overexpression drives reproducible, reversible metabolic disruption across systemic and neural compartments—anchored in mitochondrial respiration and nucleotide synthesis.

## Mitochondria in MDS are Structurally, Functionally, and Bioenergetically Impaired

Given these systemic perturbations, we next examined mitochondria directly in patient-derived cortical organoids, which recapitulate early human cortical development^33^. Untargeted metabolomics of 60-day-old MDS organoids revealed pronounced TCA cycle perturbations and substantial accumulation of multiple intermediates and related byproducts, including itaconic acid, citrate, aconitic acid, malate, succinate, fumarate, and 2HG—a canonical marker of mitochondrial dysfunction^34,35^ (**Fig. 2A**). Mass spectrometry imaging (MSI)^36^ confirmed focal accumulation of these intermediates in neuronal regions and also demonstrated that succinate abundance correlated with MECP2 expression, normalized following ASO treatment of MDS organoids (**Extended Data Fig. 4**).

**Figure 2.**
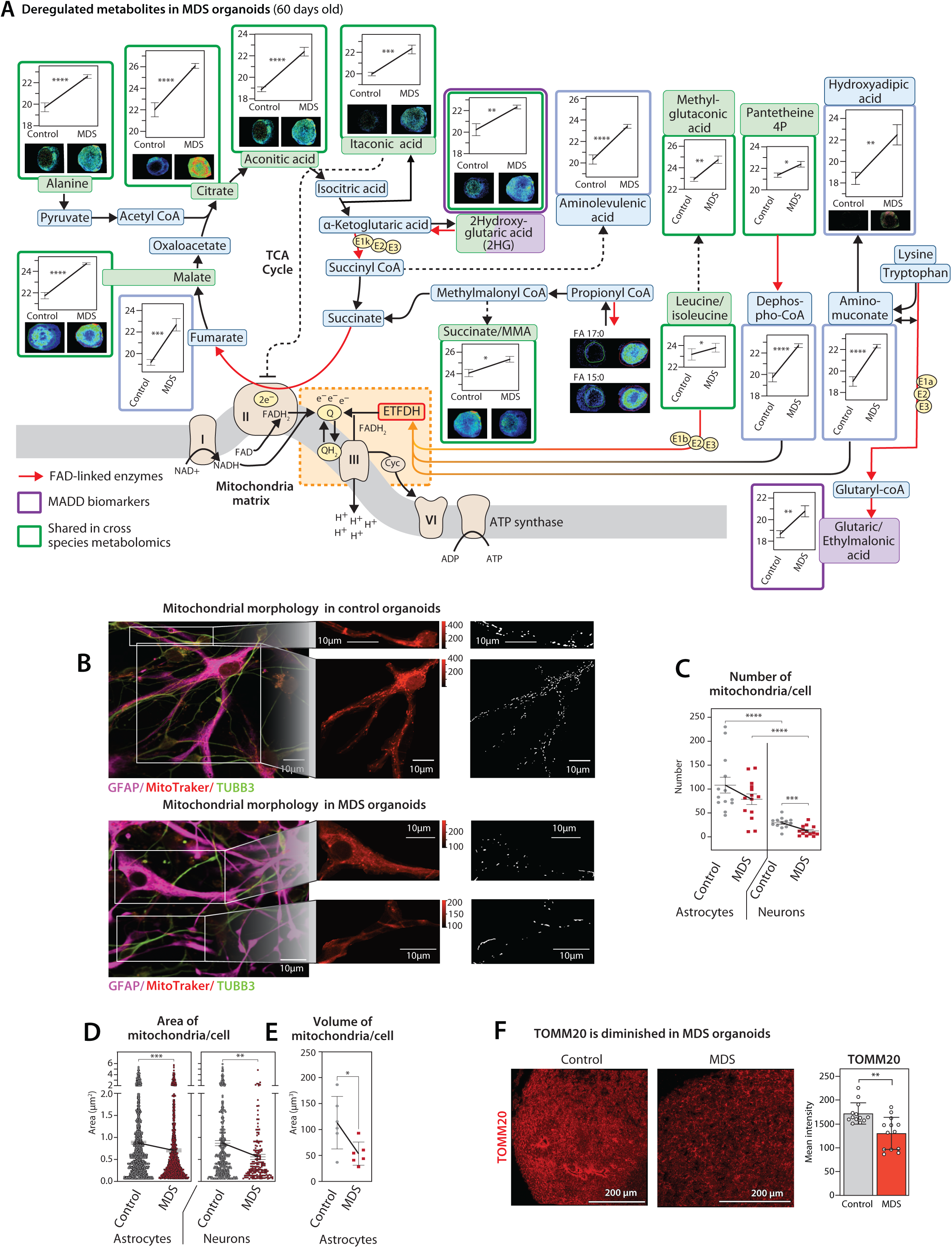
Mitochondria in MDS are structurally, functionally, and bioenergetically impaired. **A**. Metabolomics of 60-day-old MDS organoids shows significant deregulation of mitochondrial pathways, TCA cycle, and electron transport chain (ETC). Each box shows a given metabolite abundance in organoid lysates (mean ± SEM) and spatial distribution based on mass spectrometry imaging (MSI) of organoid sections. Lines connecting box-plots are for easier visualization of the differences. Boxes outlined in green are shared across species in both systemic and neurocentric metabolomics datasets. Many of the highly abundant metabolites are substrates for mitochondrial enzymes that require FAD as a cofactor (red arrows). Dotted arrows outline indirect connections between metabolites. Orange box focuses on the ETFDH enzyme that mediates transition between Complex II (CII) and CIII. Purple box indicates metabolites impaired in Multiple acyl-CoA dehydrogenase deficiency (MADD). FAD=flavin adenine dinucleotide, ETFDH=electron-transferring flavoprotein dehydrogenase, TCA=tricarboxylic acid, CoA=Coenzyme A. **B.** Confocal photomicrograph of mitochondria (MitoTracker immunostaining) in astrocytes (GFAP+) and neurons (TUBB3+) of 60-day-old organoids shows reduced mitochondrial number and a shift toward spherical morphology in MDS compared to controls in both cell types. Scale bar=10 µm. **C-E.** Quantitative analysis (mean ± SEM) of mitochondrial number (**C**), area (**D**), and volume (**E**) per cell shows overall reduction in MDS organoids compared to controls. In **C**, each dot represents a cell (control n=14, MDS n=14). In **D**, each dot corresponds to a mitochondrion identified using Mitochondria Analyzer software. In **E**, each dot corresponds to a mitochondrion, with volume quantified from z-stack images (step size 0.5µm, 10 steps total). Comparisons were done using t-test, *p<0.05, **p<0.01, ***p<0.001, ****p<0.001. **F**. Confocal photomicrograph of TOMM20 immunostaining of mitochondrial outer membrane in 60-day-old organoids shows reduced expression in MDS compared to controls. Scale bar=200µm. Bar graph shows mean intensity ±SEM (t-test, **p<0.01). Each dot is a region-of-interest (ROI); 13 ROIs were examined per organoid. All experiments on organoids used triplicates from 5-10 organoids per experiment/section, in two independent batches. Please see **Extended Data** Figures 4-6 and **Video 1** for details.

To ascertain disrupted pathways, we turned to Mitomics database, encompassing 205 human mitochondrial gene knockouts (mitoKO)^37^. Mitomics analyses revealed a significant upregulation of these same TCA metabolites (p<0.001) particularly in OXPHOS gene deletions (e.g., *TSFM, MTO1, FASTKD2, C2orf69, NDUFS3, COX6A1*) (**Extended Data Fig. 4**, **Table 20**). Strikingly, concurrent elevations of 2HG and glutaric acid—consistently observed in MDS blood and neuro datasets (**Fig. 1D**)—associated with KOs of CoQ10 biosynthesis-ETC cycle genes (*COQ8A/B, ADCK2, ETFB, CYC1*) and mitochondrial biogenesis (*FASTKD2, TIMM17B, ETHE1, GLRX5, MFF, PINK1*) and have been linked to secondary to CoQ10 deficiency in an inborn error of metabolism: Multiple acyl-CoA dehydrogenase deficiency (MADD)^38^ (**Fig 2A**, **Extended Data Table 21**). Altogether, these findings suggest that the Complex I/II-Complex III (CI/CII-CIII) junction of the electron transport chain (ETC) downstream of the TCA cycle is defective and unable to process incoming metabolites, causing high abundance of TCA intermediates.

Mitochondria in MDS organoids were also structurally and functionally impaired. They were fragmented, smaller in area, volume, and mass, with reduced branching complexity, shorter branch lengths, thinner mean branch diameters, and decreased overall width, resulting in a more spherical morphology in both neurons and astrocytes (**Fig. 2B-E; Extended Data Fig. 5, Video 1)**. Consistently, TOMM20, SDHB, and COX4 expression was markedly reduced along with diminished ATP abundance, oxygen consumption rates (OCR), and mitochondrial *ND1* and *ND2* transcripts—reflecting impaired mitochondrial biogenesis and diminished transcriptional output of the mitochondrial genome (**Fig. 2F, Extended Data Figs. 5, 6**). Collectively, these findings confirm that mitochondrial dysfunction in MDS extends beyond metabolic imbalance to encompass profound bioenergetic failure, structural disorganization, and transcriptional impairment.

## Complex III emerges as the nodal point of mitochondrial dysfunction in MDS

To delineate the molecular basis of mitochondrial dysfunction, we integrated transcriptomic, proteomic, and functional datasets. Single-cell transcriptomics revealed broad downregulation of OXPHOS genes in MDS organoids (**Fig. 3A**), while CSF and organoid proteomics pinpointed selective depletion of CIII–associated proteins (e.g., UQCRQ, UQCRB, UQCRFS1). Notably, UQCRQ emerged as a shared downregulated target across both datasets, harboring a predicted MECP2-binding domain (**Fig. 3B, C**). CUT&RUN profiling^39^ further confirmed direct MECP2 binding to promoters of UQCRQ and other CIII genes (UQCRFS1, UQCRB), all of which were downregulated in MDS organoids (**Fig. 3D; Extended Data Fig. 6**). Notably, UQCRQ expression was restored by ASO (**Fig. 3E-F**).

**Figure 3.**
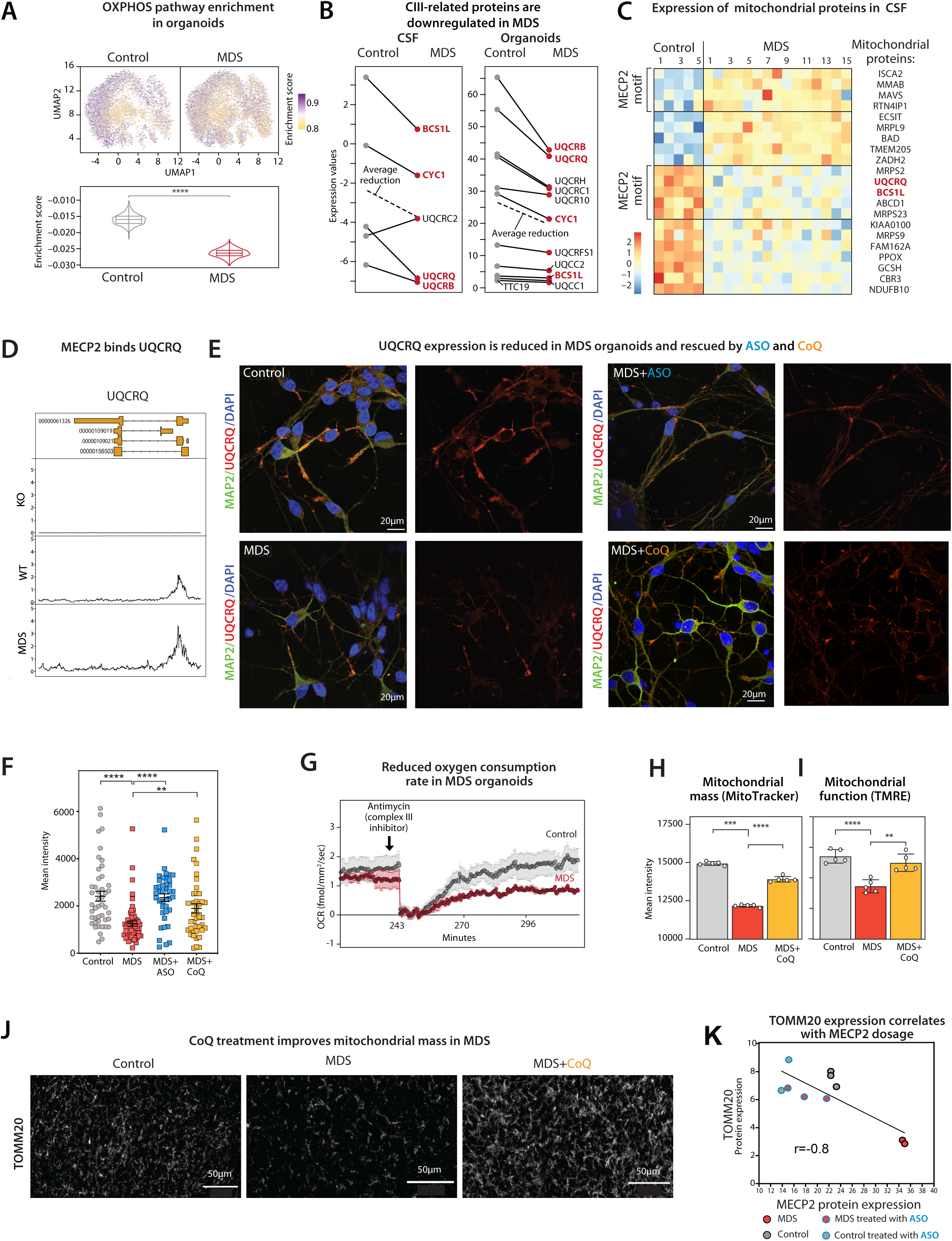
Complex III deficiency drives mitochondrial dysfunction in MDS. **A**. Single cell-RNAseq (scRNAseq) of MDS organoids at 45-day-old shows marked downregulation of the oxidative phosphorylation (OXPHOS) pathway, visualized by UMAP and violin plots (Wilcoxon rank-sum test, ****p<0.001). **B.** Proteomics of CSF and cortical organoids from MDS patients reveals significant (p<0.05) decrease in multiple Complex III (CIII) proteins (mean reduction indicated by a dotted line), consistent with ETC dysfunction. Each dot is the average expression of the targeted proteins. Connecting lines are for easier visualization of the differences. **C.** CSF proteome heatmap shows reduced mitochondrial proteins (UQCRQ, BCS1L, GCSH, NDUFB10) in MDS, with UQCRQ and BCS1L predicted to contain MECP2-binding motifs. Each column is a sample, each row a mitochondrial protein. Red letters denote proteins shared between CSF and organoids. **D.** CUT&RUN analysis of MDS and WT mice shows MECP2 binding at the UQCRQ locus, which was increased in MDS and completely absent in MECP2 knockout (KO) mice (data from^41^). The top box corresponds to mouse gene identifier. **E–F** Confocal images of UQCRQ in MAP2⁺ neurons show reduced expression in MDS, which was restored to control levels following ASO (2.5 μM) or CoQ10 (5 μM) treatment. **F** Quantification of MAP2⁺ ROIs revealed a significant increase in UQCRQ intensity after both treatments (each dot represents one ROI; 10–15 neurons per condition from three biological replicates were analyzed). Scale bar, 20 μm.**G.** Oxygen consumption rate (OCR) measured by Resipher is reduced in MDS organoids at baseline and recovery after antimycin A (CIII inhibitor) is slower than in controls (t-test, ***p<0.001). **H-I.** Bar graphs (mean±SEM) show that CoQ_10_ increases mitochondrial mass (Mitotracker flow cytometry, **H**) and improves mitochondrial membrane potential (Tetramethylrhodamine ethyl ester (TMRE) flow cytometry; **I)** in MDS organoids. Each dot is a sample from 3-4 60-day-old disaggregated organoids done in triplicate, in three independent batches. t-test, ***p<0.001, ****p<0.001. **J**. Confocal photomicrographs of TOMM20 mitochondrial inner membrane immunostaining shows that CoQ_10_ supplementation increases mitochondrial mass in 60-day-old MDS organoid. Scale bar=50µm. **K.** Protein expression of TOMM20 inversely correlates with MECP2 dosage (Pearson correlation, r=-0.8, **p<0.05). Data points are color-coded by condition: MDS (red), controls (gray), MDS treated with ASO (red with blue edge), controls treated with ASO (gray with blue edge). MECP2 overexpression is associated with reduced TOMM20 levels, whereas MECP2 knockdown via ASO restores (in MDS) or increases (in controls) TOMM20 expression. Please see **Extended Data** Figure 6 and **Tables 24-25** for details.

Functionally, MDS organoids were hypersensitive to the CIII inhibitor antimycin, exhibiting a delayed recovery of mitochondrial respiration compared to controls (**Fig. 3G**) and further supporting impaired CIII resilience and reduced bioenergetic flexibility with MECP2 overexpression. Given that oxidized CoQ_10_—an essential electron carrier and biochemical marker of CIII deficiency—was significantly reduced in MDS organoids (**Extended Data Fig. 6**), we next tested whether CoQ_10_ supplementation could rescue mitochondrial defects. Remarkably, CoQ_10_ treatment reinstated UQCRQ expression (**Fig. 3E-F**) and improved mitochondrial mass and membrane potential (**Fig. 3H-J**), establishing a causal link between CoQ_10_-dependent CIII activity and MECP2-driven mitochondrial dysfunction.

Interestingly, TOMM20 levels correlate with CIII and OXPHOS activity in human brain tissue^40^. TOMM20 expression was restored by ASO (**Fig. 3K**), suggesting that improved mitochondrial organization arises secondary to CIII recovery.

Mechanistically, CIII impairment limits ubiquinol oxidation^41–44^, disrupting the CoQ-cycle and slowing electron flow through the ETC. This causes accumulation of reduced electron carriers (NADH, FADH_2_)^45^ and feedback inhibition of multiple NAD⁺- and FAD-dependent TCA dehydrogenases, stalling TCA flux^46,47^. Consistently, MDS organoids exhibited elevated NADH, decreased NAD⁺ and FAD (**Extended Data Figs. 3, 6**), and accumulated TCA intermediates— succinate, fumarate, malate, and aspartate—indicating congestion at the CI/II–III junction. This metabolic configuration reflects a state of “reductive pressure”, further supported by increased polyamine-derived (spermidine) metabolites, byproducts of excess in reduced equivalents^48^ (**Extended Data Figs. 3**). Importantly, several FAD-dependent enzymes, including protoporphyrinogen oxidase (PPOX), were downregulated (**Extended Data Fig. 7)**. PPOX is part of the mitochondrial heme pathway^49^ and its loss likely limits heme availability for CIII assembly, establishing a vicious cycle where CIII dysfunction and impaired heme synthesis amplify redox imbalance. Moreover, as FAD is a cofactor also for L-2HG dehydrogenase that converts 2HG to α-ketoglutarate^50^, reduced FAD may contribute to the elevated 2HG accumulation we consistently observed across all MDS datasets (**Fig. 2A**). Altogether, these data position CIII as the critical bottleneck linking redox imbalance to global metabolic collapse in MDS.

To examine whether the MECP2–CIII relationship is bidirectional, we examined published datasets on Rett syndrome. Independent transcriptomic and proteomic studies consistently reported opposite trend: upregulation of CIII components—including UQCRQ, UQCRFS1, UQCRC1, and UQCRH—accompanied by combined CI/III alterations and an elevated FAD/NADH ratio in both brain and peripheral tissues^25,51–55^. Consistently, in Rett astrocytes, TCA intermediates—citrate and cis-aconitate—are diminished^56^. This inverse pattern reinforces a dosage-sensitive, bidirectional regulation of CIII by MECP2, positioning mitochondrial bioenergetic imbalance as a unifying mechanism across the MECP2 dosage spectrum.

## CIII dysfunction drives oxidative stress, DNA damage, and purine biosynthesis

Importantly, CIII dysfunction triggers reactive oxygen species (ROS) generation^57^ and oxidative DNA damage^38,58,59^, including double-strand breaks^60^. Consistently, in MDS organoids we observed elevated ROS and 8-hydroxy-guanosine—a marker of oxidative nucleic acid damage—accompanied by oxidized glutathione (GSSG) accumulation and diminished reduced glutathione (GSH) (**Fig. 4A–C, E; Extended Data Fig. 7**), indicating a compromised redox buffering capacity. Pyroglutamic acid—a γ-glutamyl cycle byproduct that accumulates when GSH synthesis is impaired or demand is excessive^61^—was increased in MDS organoids (**Fig. 4D**) and cross-species blood (**Extended Data Fig. 2**), supporting sustained oxidative stress and aligning with prior clinical reports of elevated oxidative stress markers in MDS^62^.

**Figure 4.**
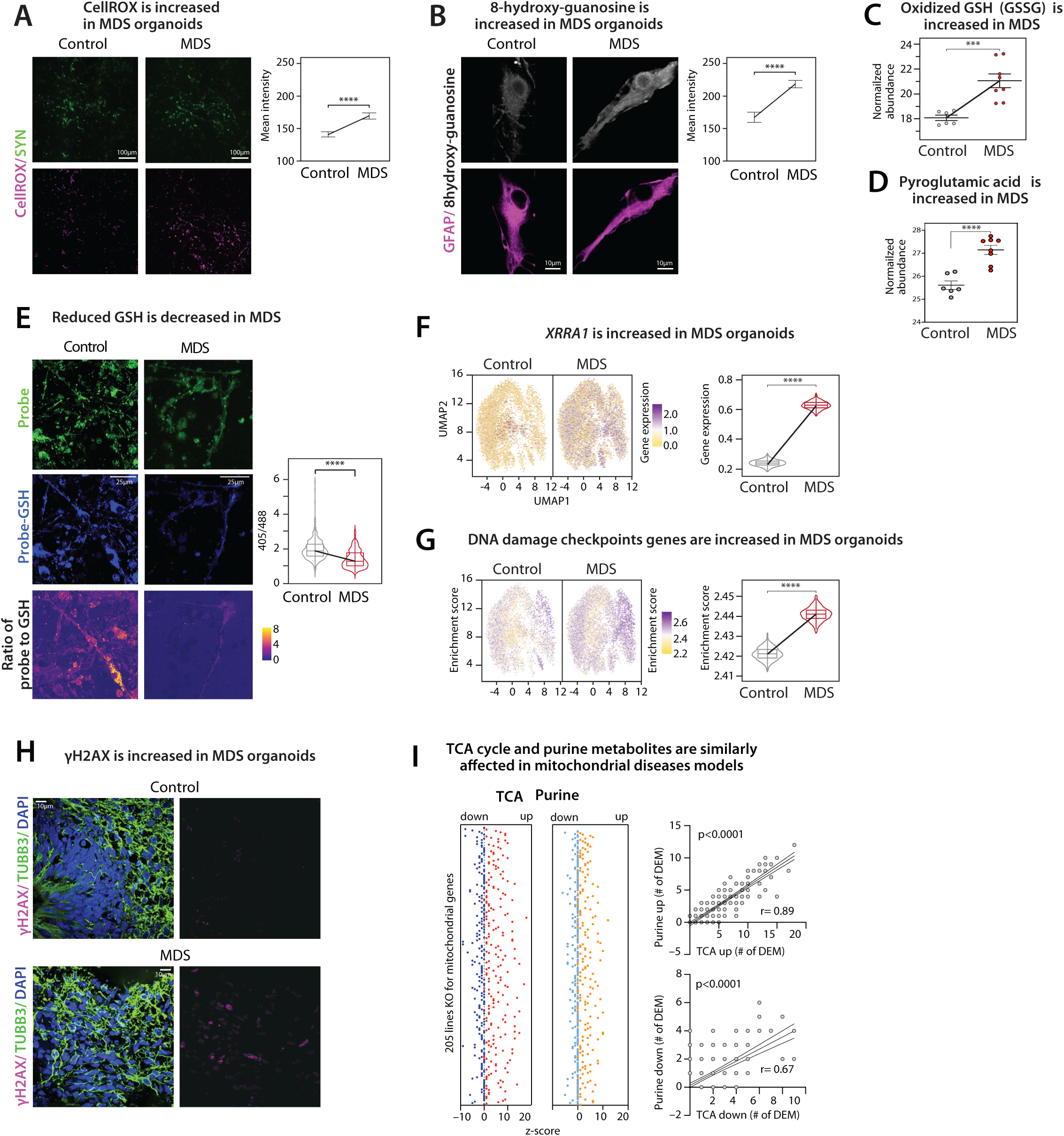
Elevated oxidative stress in MDS leads to DNA damage and metabolic pathway alterations. **A.** Confocal photomicrographs show increased CellROX fluorescence mean intensity in Syn1^+^ neurons in MDS organoids, indicating elevated oxidative stress compared to controls. Scale bar=100 µm. Quantitative analysis (mean±SEM; t-test, ****p<0.001) is shown on the right. **B.** Confocal photomicrographs show significantly increased 8-hydroxy-guanosine, a marker of the oxidative damage, in GFAP^+^ astrocytes in MDS organoids. Scale bar=10 µm. Quantitative analysis (mean±SEM; t-test, ****p<0.001) is shown on the right. **C.** Oxidized glutathione (or GSSG) is increased in MDS organoid lysates as measured by LC-MS. Each dot is a sample from lysates of twenty 60-day-old organoids (four biological replicates, two independent batches; t-test, ***p<0.001). **D.** Pyroglutamic acid, a byproduct of glutathione (GSH) metabolism and marker of GSH deficiency, is significantly increased in MDS organoid lysates as measured by LC-MS. Each dot represents a sample derived from lysates of twenty 60-day-old organoids (four biological replicates, two independent experimental batches; *t*-test, \*\**p* < 0.001). **E.** Multi-photon photomicrographs show decreased levels of reduced glutathione (GSH) in MDS. Scale bar=25µm. Violin plots show quantitative analysis of GSH levels (ratio of fluorescence intensity of the probe measuring free GSH and the probe bound to GSH)^110^, t-test, ****p<0.001. **F.** scRNAseq shows upregulation of *XRRA1*, a gene associated with cellular responses to oxidative stress, in MDS organoids (*XRRA1* is upregulated also in bulk RNAseq data at both 30- and 60-day-old MDS organoids; see **Extended Data Fig. 7**) (Wilcoxon rank-sum test, ****p<0.001)). **G.** DNA damage checkpoint genes are increased in MDS organoid scRNAseq compared to controls (Wilcoxon rank-sum test, ****p<0.001). Pathway enrichment score violin plots are shown on the right. List of genes associated with DNA damage checkpoints are detailed in **Methods**. **H.** Confocal photomicrographs show increased γH2AX staining (DNA damage marker) in the MDS organoid. Scale bar=10µm. **I.** In the Mitomics dataset (205 cell lines with knockouts (KOs) of genes encoding mitochondrial proteins), the abundance of TCA cycle metabolites strongly correlates with that of purine-related metabolites (Pearson correlation, *p* < 0.0001). This positive association highlights the interdependence between energy metabolism and nucleotide biosynthesis. DEM = differentially expressed metabolites. Please see **Extended Data** Figures 7 for details.

Persistent oxidative stress in MDS organoids was accompanied by robust activation of the DNA damage response, marked by increased expression of *XRRA1*^63,64^, *CCND1, BRCA1,* and key regulators of the DNA damage signaling cascade, including TP53 and ATR, along the pronounced γH2AX accumulation (**Fig. 4F–H; Extended Data Fig. 7,8**). To exclude the possibility that elevated γH2AX reflected increased replication activity, we assessed EdU incorporation after one week of treatment. We observed a significant reduction in EdU^+^ nuclei in MDS organoids, indicating decreased DNA synthesis, alongside increased γH2AX foci— demonstrating that DNA damage arises from oxidative and replicative stress rather than hyperreplication (**Extended Data Fig. 7**).

Activation of DNA damage response pathways elevates purine demand, as purines serve as ROS sensors^65^ and early DNA damage markers^66^. Elevated ROS boosts purine metabolism^67^ and shifts glucose metabolism toward the pentose phosphate pathway to supply nucleotides for DNA repair^68^. As mitochondria regulate nucleotide biosynthesis essential for DNA repair via coordinated substrate generation^69–71^, we examined whether mitochondrial dysfunction in MDS is linked to purine metabolism. In Mitomics datasets, we observed strong correlations between TCA cycle intermediates and purine metabolite abundance across mitoKO lines (**Fig. 4I**). This association was particularly pronounced in *CYC1*-deficient cells—a core CIII component— where metabolic signatures closely mirrored those observed in MDS (**Extended Data Fig. 8**).

*De novo* purine biosynthesis is compartmentalized within purinosomes—cytoplasmic, multienzyme condensates that assemble in proximity to mitochondria to exploit locally enriched metabolic precursors and coordinate nucleotide synthesis with cellular energy and redox balance^72^. This spatial coupling enhances pathway efficiency and minimizes loss of intermediates^69^. In MDS organoids, key purinosome enzymes were markedly increased in both radial glia (SOX2⁺) and neuronal (TUBB3⁺) regions (**Fig. 5A**) and the overall number (controls=125; MDS=242), area, and fluorescence intensity of purinosome puncta were significantly elevated in MDS, indicating enhanced purinosome assembly (**Fig. 5B**; **Extended Data Figs. 9**). Consistent with these expression signatures, purine metabolites—including adenine, adenosine, AMP, ADP, cyclic ADP ribose, ADP ribose, deoxyadenosine, adenylosuccinic acid, guanosine, guanine, GMP, inosine, and IMP—were significantly elevated in MDS organoids (**Fig. 5D**). Fumarate and aspartate—two metabolites that mediate mitochondria–purinosome coupling—were also increased (**Fig. 5E**), indicating enhanced flux through this metabolic interface. These data were supported by both transcriptomics and proteomics data, which showed upregulation of ADSS2, ENPP2, ENPP3, GART, and PFAS— the core enzymes of the *de novo* purine biosynthesis (**Extended Data Fig. 8, 9**). Activation of the pentose phosphate pathway further supported a metabolic shift from glycolysis toward nucleotide synthesis (**Extended Data Fig. 8**). Interestingly, PFAS expression was elevated in MDS organoids and normalized by ASO (**Fig. 5C**; **Extended Data Fig. 9**), which also restored adenine abundance, confirming MECP2-dependent regulation of purine-related enzymes and metabolites (**Fig. 5G**).

**Figure 5.**
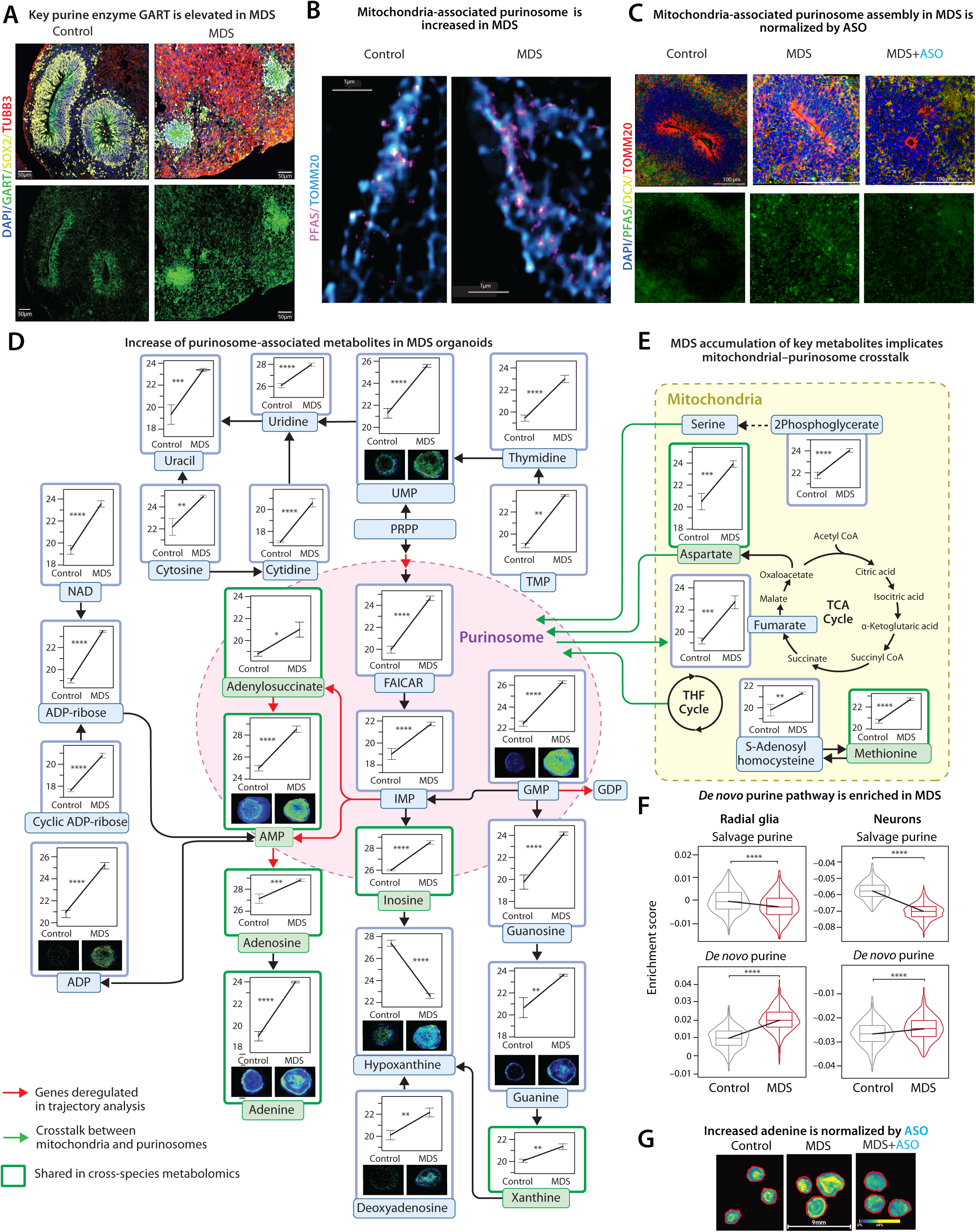
Elevated purine metabolites in MDS accumulate in purinosomes attached to mitochondria. **A.** Confocal photomicrographs show increased expression of GART, a critical enzyme in purine biosynthesis, in rosettes (SOX2^+^) and neurons (TUBB3^+^) in the MDS organoid. Scale bar=50µm. **B.** Ultra-resolution Stimulated Emission Depletion (STED) microscopy shows increased proximity of PFAS^+^ purinosomes and TOMM20^+^ mitochondria. In MDS, there is an increase in purinosome area, puncta number, and fluorescence intensity compared to controls. Scale bar=1µm. **C**. Confocal photomicrographs show increased expression of PFAS in MDS, normalized after 2.5 µM ASO treatment. Scale bar=100µm. **D.** Metabolomics of 60-day-old MDS organoids shows significant upregulation of purines. Each box shows a given metabolite abundance in organoid lysates (mean±SEM) and spatial distribution based on mass spectrometry imaging (MSI) of organoid sections. Lines connecting box-plots are for easier visualization of the differences. Boxes outlined in green are shared across species in both systemic and neurocentric metabolomics datasets. Black arrows show direct relationships between given metabolites. Red arrows show pathways with deregulated enzymes based on the sc-RNAseq data on the same organoids. Purines are assembled into purinosomes indicated by the pink circle (t-test, **p<0.01, ***p<0.001, ****p<0.001). **E.** Mitochondrial metabolites such as aspartate and fumarate (essential for both TCA cycle and *de novo* purine synthesis) are elevated in MDS. Each box shows a given metabolite abundance in organoid lysates (mean±SEM). Lines connecting box-plots are for easier visualization of the differences. Boxes outlined in green are shared across species in both systemic and neurocentric metabolomics datasets. Black solid arrows show direct relationships between given metabolites. Dotted arrow shows indirect connection between metabolites. Green arrows show direct crosstalk metabolites between mitochondria and purinosomes. All metabolites and pathways within mitochondria are within the yellow box. t-test, **p<0.01, ***p<0.001, ****p<0.001. THF= tetrahydrofolate cycle (carrier for one-carbon units for purine synthesis), FAICAR= 5-formamidoimidazole-4-carboxamide ribonucleotide. **F.** Pathway analysis of scRNAseq shows significant enrichment of *de novo* purine synthesis pathway in radial glia and neurons compared to the salvage pathway, which is diminished (Wilcoxon rank-sum test, **p<0.01, *\*\*\*\**p<0.001). **G.** MSI distribution of adenine shows increased abundance in MDS compared to controls, normalized following ASO treatment (2.5 µM). Please see **Extended Data** Figures 8–9 for detailed analyses.

Together, these data establish that CIII dysfunction induces purinosome condensation, coupling mitochondrial collapse to hyperactive nucleotide synthesis. This spatial and functional coupling generates a maladaptive feedback loop—mitochondrial stress increases ROS and DNA damage, which further amplifies purine biosynthesis to sustain repair, perpetuating redox imbalance and metabolic overload.

## Mitochondrial-purinosome dysfunction disrupts neurogenesis

Having identified the mitochondria–purinosome axis as central to MDS pathology, we next examined its developmental consequences. MDS organoids displayed a marked reduction in the number, wall thickness, and overall area of ventricular-like rosettes, consistent with impairment of radial glia—the primary stem cells of the developing brain (**Fig. 6A, B**). Single-cell transcriptomics confirmed downregulation of progenitor markers (*PAX6, MKI67, TOP2A*) and diminished phospho-vimentin staining and EdU incorporation indicated reduced mitotic activity (**Fig. 6C, D; Extended Data Figs. 7, 10**). Metabolic data further reinforced these findings, revealing disruption of the glutamate-glutamine cycle—a critical pathway linking redox and neuroprogenitor homeostasis—across transcriptome and metabolome datasets (**Extended Data Fig. 10**). Radial glia, reliant on glycolysis yet dependent on mitochondrial one-carbon and nucleotide flux for replication, appeared uniquely vulnerable.

**Figure 6.**
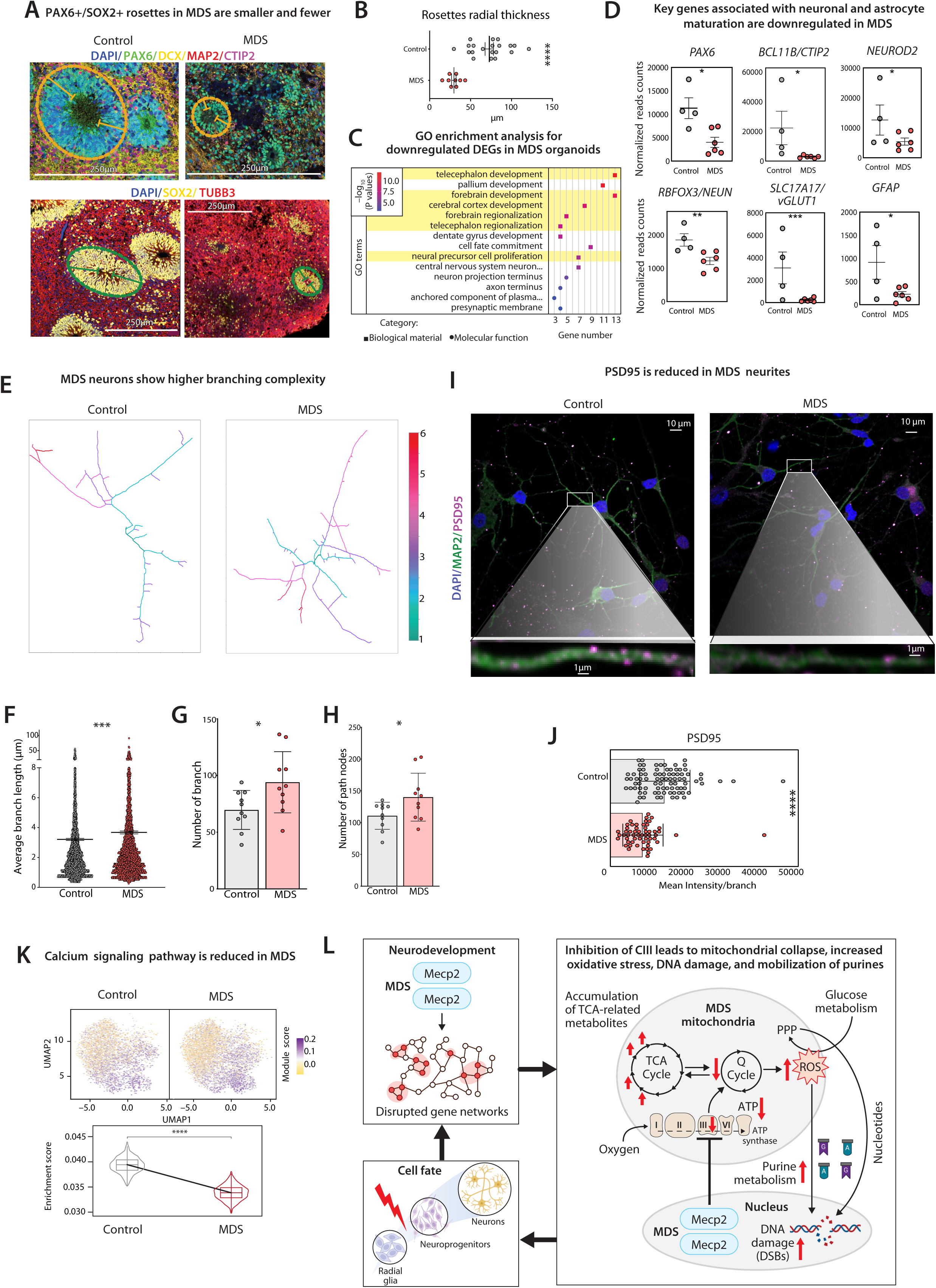
Mitochondrial-purinosome dysfunction disrupts neurogenesis. **A.** Confocal micrographs show PAX6^+^ (upper panel) and SOX2^+^ (lower panel) rosettes, immature DCX^+^ neurons, more mature MAP2^+^ and TUBB3^+^ neurons, and layer II CTIP2^+^ neurons. Rosettes are outlined with dotted lines and measured diameters with solid lines. Scale bar=250 µm. **B**. Quantitative analysis of rosette wall thickness (outer–inner edge distance) in 60-day organoids (mean ± SEM; *t*-test, ****p<0.0001).. **C.** GO enrichment analysis of downregulated differentially expressed genes (DEGs) in 60-day-old MDS organoids shows pathways associated with neural cell fate determination during early cortical development (highlighted in yellow). **D.** Key genes associated with radial glia and neuroprogenitor identity, neuronal differentiation, and astrocyte maturation are significantly downregulated in the transcriptome of 60-day-old MDS organoids (mean ± SEM; *t*-test, *p* < 0.05, \**p* < 0.01, \*\**p* < 0.001). **E-H.** 2D reconstruction of neurons (Neurite Tracer plugin in Fiji) illustrates branch complexity (color-coded from 1 to 6) and increased branching in MDS neurons compared to controls (**E**). Quantitative analyses show increased average branch length (**F**), higher number of branches (**G**), and greater number of nodes (**H**) in MDS neurons. For this analysis, MAP2⁺ neurons were selected and at least ten images per condition were analyzed and normalized to the number of neurons (63× magnification). **I-J.** Confocal photomicrographs show a significant reduction in the number and fluorescence intensity of the postsynaptic protein PSD95 within dendrites of MAP2⁺ MDS neurons (**I**). Scale bars=10µm and 1µm. PSD95 puncta quantification was done in Fiji; mean intensity was measured per object within the PSD95 mask. (**J**). **K.** Calcium signaling pathway is downregulated in scRNAseq MDS organoid transcriptome in neuronal clusters compared to controls. Pathway enrichment score violin plots are shown (Wilcoxon rank-sum test, ***p<0.001). **L.** Proposed model outlines the effect of MECP2 overexpression on gene networks and metabolism, where direct inhibition of CIII leads to TCA cycle metabolite accumulation, increased oxidative stress, DNA damage, and enhanced purine metabolism, which together contribute to radial glia and neuroprogenitor cell fate derailment and delayed neuronal maturation. PPP=Pentose Phosphate Pathway. DSB=Double strand break. See **Extended Data Fig. 10** for details.

Further, the number of newborn EdU⁺/DCX⁺ neurons was significantly reduced (**Extended Data Fig. 10**), along with broad downregulation of cortical development genes (**Fig. 6D**). MDS neurons showed significantly increased branch length and a higher number of nodes, consistent with previous reports^16,73,74^ (**Fig. 6 E-H**). Yet, PSD95 expression was significantly reduced — (**Fig. 6I, J**), accompanied by decreased calcium activity (**Fig. 6K**)—critical for neuronal excitability and network activity^75^—indicating hyperextended but hypofunctional maturation.

Interestingly, purine supplementation of the culture media enhances neurite extension and branching complexity in both control and diseased neurons^76,77^, consistent with observations that extracellular purines and their receptors significantly influence neurite outgrowth and neuronal morphology^78^. Conversely, purine deficiency leads to shorter neurites, reduced branching, and defective synaptogenesis^79^. In line with these observations, MDS organoids exhibited increased purine-related metabolites and expression of specific purinergic receptors, including ADORA and P2RX3 (**Extended Data Fig. 9**), suggesting an adaptive response to heightened purine signaling.

Thus, *MECP2*-driven mitochondria-purinosome dysfunction extends beyond bioenergetic failure to directly alter neurogenesis in the developing brain. By directly impairing radial glia transcriptional programs and proliferation and delaying neuronal maturation, excess *MECP2* distorts cortical trajectories. Although direct MECP2 binding to radial glia genes likely contributes^80^, our multiomics and functional evidence firmly establishes that the dominant driver is metabolic disintegration: a failure of mitochondria to sustain biosynthetic flux for genome replication and repair, derailing neuroprogenitor dynamics and cortical development in MDS.

## Mechanistic model: the mitochondria-purinosome axis as a developmental control point

Together, our findings delineate a pathogenic cascade in which MECP2 overexpression disrupts mitochondrial CIII, triggering TCA-intermediates accumulation, redox collapse, oxidative stress, DNA damage, and compensatory *de novo* purine biosynthesis (**Fig. 6L**). The resulting hyper-condensation of purinosomes around damaged mitochondria forms a self-reinforcing feedback loop that amplifies metabolic stress and genomic instability. This cascade ultimately compromises radial glia proliferation, delays neuronal differentiation, and perturbs neurodevelopmental trajectories of cortical circuits. In essence, mitochondrial failure in MDS is not merely a secondary byproduct but a primary determinant of neurodevelopmental derailment. The mitochondria-purinosome crosstalk thus represents a previously unrecognized metabolic-genomic interface governing neuroprogenitor fate.

## DISCUSSION

By integrating multiomics, functional, and morphological analyses across patients, organoids, and mouse models, we reveal how *MECP2* overexpression reconfigures mitochondrial metabolism and nucleotide balance. CIII emerges as the molecular fulcrum translating gene dosage imbalance into systemic metabolic and developmental pathology.

The brain’s unique reliance on oxidative metabolism renders it acutely vulnerable to mitochondrial disruption^81^. Yet, in radial glia—where glycolysis predominates^82^—mitochondria appear to fulfill a biosynthetic role, channeling one-carbon and nucleotide precursors to support DNA synthesis and genome integrity during cell division. This aligns with growing evidence that metabolic states define stem and progenitor cell fate across diverse systems^83^, but our work extends this principle to a human neurodevelopmental disorder *in vivo*. Our data suggest that the mitochondria-purinosome coupling functions as a developmental metabolic switch, ensuring that proliferating progenitors maintain DNA integrity while differentiating neurons transition to energy-intensive OXPHOS states.

Disruption of this coupling through CIII deficiency therefore impairs both bioenergetic and biosynthetic integrity, explaining why mitochondrial dysfunction manifests so prominently in early neurodevelopment. CIII lies at a critical juncture in the ETC, funneling electrons from CI and II to C IV^84–86^. Its deficiency leads to upstream accumulation of reduced intermediates, redox imbalance, and impaired ATP generation^71–78^, all of which we observed in MDS models. The consistent metabolic, proteomic, transcriptomic, and functional signatures across organoids, mouse tissues, and patient biofluids strongly support CIII as the mechanistic driver of the observed pathology. The partial rescue of metabolic and purine signatures by ASO demonstrates both causality and clinical tractability of these pathways, unifying diverse metabolic and transcriptional findings across MECP2 disorders: Rett syndrome exhibits the reciprocal upregulation of CIII components, highlighting a dosage-sensitive control of mitochondrial homeostasis.

A striking observation was the persistent activation of *de novo* purine biosynthesis across human, organoid, and mouse datasets. The purinosome, a transient multi-enzyme complex, colocalizes with mitochondria to optimize nucleotide flux for DNA replication and repair^65–68^. In MDS, this spatial and functional coupling was disrupted, perpetuating metabolite accumulation, redox imbalance, with broad consequences for genome integrity and progenitor function. While purinosome biology has been primarily defined in cancer^29,96^, and more recently in inborn errors of metabolism involving *de novo* purine defiencies^99^, our results highlight its relevance for neurodevelopmental disorders and position it as a potential therapeutic target.

Although MDS is rare, its mechanistic lessons are broadly relevant as it provides a powerful lens to interrogate how subtle perturbations in gene dosage can cascade into systemic metabolic derangements. Recent studies have discussed mitochondria–purine coupling as a critical metabolic axis^98^, which our work now implicates in neurodevelopmental pathology. The discovery that mitochondrial-purine uncoupling is a central pathogenic mechanism not only advances our understanding of MDS but also illuminates shared vulnerabilities across conditions where mitochondrial dysfunction intersects with neurodevelopment. Indeed, brain MRI abnormalities observed in MDS patients—including corpus callosum dysgenesis, delayed myelination, ventricular enlargement, and reduced white matter^100^—closely resemble those seen in primary mitochondrial disorders^101^. This striking overlap suggests a shared pathogenic mechanism, where MECP2 dysfunction may lead to mitochondrial impairment, ultimately disrupting normal neural development. Our findings support this link, highlighting mitochondrial-purine uncoupling as a likely generalizable mechanism underlying structural brain changes seen in MDS as well as in other X-linked intellectual disability syndromes and primary mitochondrial disorders. This conceptual framework positions mitochondria not simply as energy factories but as architects of genomic and developmental stability—a view that opens new avenues for therapeutic intervention through metabolic restoration.

In sum, by delineating a causal chain from *MECP2* overexpression to CIII dysfunction, redox imbalance, and purine hyperactivation, and disrupted neurogenesis, this work provides a unifying model of how transcriptional misregulation reconfigures cellular metabolism to derail brain development. By situating mitochondria as the gatekeepers of both bioenergetic and biosynthetic integrity, our findings bridge a longstanding gap between metabolic dysfunction and neurodevelopmental outcomes. Reestablishing mitochondrial-purine coupling thus emerges as a mechanistic and translational goal for restoring metabolic and developmental equilibrium in gene dosage-sensitive disorders.

## ACKNOWLEDGEMENTS

This work was supported in part by grants from the Eunice Kennedy Shriver National Institute of Child Health & Human Development of the National Institutes of Health (P50HD103555 to M.M.-S. and core facilities), and the Genomic and RNA Profiling Core at Baylor College of Medicine. In addition, this work was supported in part by the Ionis Pharmaceuticals, grants from the National Institutes of Health (R01MH130356 to M.M-S.), Autism Speaks (G.C.), and Cynthia and Antony Petrello Endowment (M.M.S.). D.P. is supported by the Doris Duke Charitable Foundation (#2023-0235) and NINDS 1K23 NS125126-01A1. The content of this paper is solely the responsibility of the authors and does not necessarily represent the official views of the National Institutes of Health.

## AUTHOR CONTRIBUTION

G.C. designed and performed the experiments, analyzed and interpreted the data, and wrote the manuscript. G.Q., H.C., and Z.L. performed bioinformatic analyses, and contributed to manuscript writing. J.H., Y.L., J.M., and A.S. designed and conducted mouse experiments. S.O., S.S., T.C., J.S., and G.T. assisted with cortical organoid generation, maintenance, sample preparation, and data collection. L.P. coordinated and collected human samples. D.R.I., X.Q., S.K., and F.L. performed metabolomic and mass spectrometry imaging experiments, including data acquisition and interpretation. B.S. and D.P. clinically evaluated MDS patients and coordinated sample collection. S.A. conducted flow cytometry experiments. P.J.-N. and C.C. participated in mouse and human data collection and interpretation, and H.Z. provided intellectual guidance. M.M.S. designed and supervised all experiments, analyzed and interpreted the data, provided financial support, and co-wrote the manuscript. All authors discussed the results and approved the final version of the manuscript.

## DECLARATION OF INTEREST

J.H., Y.L., J.M., A.S., C.C. and PJ.-N. are paid employees of Ionis Pharmaceuticals. D.P. provides consulting service for Ionis Pharmaceuticals, M2DS Therapeutics and Acadia Pharmaceuticals.

## LEGEND TO EXTENDED DATA FIGURES

**Extended Data Fig. 1. A.** The study cohort consisted of 30 patients with MDS (age range 1.9-31 years, mean 10.48 years). From these patients, we collected plasma samples (including repeat sampling from 10 patients), cerebrospinal fluid (CSF; sampled twice per patient), and fibroblasts from 4 patients for iPSC generation. **B.** Genomic coordinates (Genomic assembly Hg19) of MDS patients enrolled (n=30) in the study. Two individuals are twins. In red are highlighted individuals from whom iPSCs were derived to generate brain organoids. **C.** Quality control of control iPSC lines (GM04545, GM23815, GM01888) included alkaline phosphatase staining, pluripotency score assessment (Thermo Fisher), and KaryoStat™ analysis. All three lines showed robust alkaline phosphatase activity and high pluripotency scores, consistent with a fully reprogrammed pluripotent state. KaryoStat™ analysis confirmed a male (XY) genotype and revealed no detectable chromosomal abnormalities. The MDS iPSC lines have been previously characterized and described^102^. **D***. Immunostaining of brain organoids.* Immunostaining of brain organoids from control and MDS lines at 60 days of age confirms increased MECP2 expression in MDS organoids in both neural progenitor regions (SOX2^+^) and neuronal compartments (MAP2^+^), with particularly higher abundance in the latter. **E***. Single-cell RNA-seq analysis.* Single-cell transcriptomic profiling in 60-day-old organoids revealed altered cluster distribution in MDS compared to controls. *MECP2* expression is elevated across multiple clusters (radial glia included), with the highest expression in neuron-related clusters. **F.** The mouse blood cohort included 53 MDS mice (8–12 weeks old), whose 35 were sampled under basal conditions and 18 were treated with ASO prior to blood collection. **G.** The mouse cortex cohort included 14 MDS mice (8–12 weeks old), with 8 sampled under basal conditions and 6 treated with ASO before cortex collection **H-I.** *MECP2 expression in brain organoids.* Protein (**H**) and transcript (**I**) analyses revealed significantly elevated MECP2 levels in MDS-derived brain organoids at both day 30 and day 60. **J.** Brain organoids were treated with antisense oligonucleotides (ASOs) at two doses, 2.5 μM and 5 μM, with two administrations given every other day at 51 and 53 days of age, alongside an ASO control. Samples were collected two weeks after treatment. The 2.5 μM dose at two weeks post-treatment was selected for downstream analyses, as it represented the lowest concentration that produced a significant reduction in MECP2 expression compared with untreated and scrambled controls. **K–L.** Quantitative analysis of *Mecp2* mRNA levels in the MDS mouse model. *Mecp2* expression in MDS mice is shown as a percentage relative to WT, confirming overexpression in the disease model (**K**). In parallel, *Mecp2* levels in ASO-treated MDS mice are expressed relative to PBS-treated MDS controls, demonstrating a significant reduction of *Mecp2* following treatment (**L**).

**Extended Data Fig. 2.** *Analysis of blood metabolomics datasets in human and mouse models of MDS.* **A.** *Human blood dataset analysis.* Variable Importance in Projection (VIP) scores highlight metabolites distinguishing MDS patients from controls. A volcano plot illustrates significantly altered metabolites (logFC > |0.6| and *p* < 0.05). Key metabolites involved in redox balance and mitochondrial function are labeled, including 2-methylcitric acid, decanoyl carnitine, and dodecenoylcarnitine, oxidized glutathione and ubiquinone. Heatmap and hierarchical clustering shows clear discrimination between MDS and control samples. A subset of differential metabolites is labeled; the complete list is available in **Extended Data Table 2**. **B**. Validation of key metabolites (adenine, adenosine monophosphate, 2-hydroxyglutarate) from MDS and controls samples’ cohort, using internal standard (IS) is shown. **C.** *Mouse blood dataset analysis*. VIP scores and volcano plot for MDS model mice vs wild-type (WT) controls reveal significantly altered metabolites (logFC > |0.6| and *p* < 0.05), including reduced nicotinamide riboside, hydroxyadipic acid, reduced glutathione, and decanoyl carnitine. Notably, decanoyl carnitine is a shared marker between mouse and human datasets. Heatmap and clustering shows strong group separation between MDS and WT samples. A list of significantly altered metabolites is presented (full list in **Extended Data Table 2). D.** Targeted and complementary Global Metabolomics and Pathway Screening (GMAP) analyses revealed overlapping distributions for several detected metabolites, including adenine, malate, and 2-hydroxyglutarate (2HG), showing consistent increases in z-score distributions across the systemic dataset and confirming findings observed in patient-derived samples. **E** Plots of 2-hydroxyglutarate and citrate in patients (black) and controls (red). Patients showed modest negative correlations with age, which were weak and absent in controls; therefore, age was not used as a covariate (p > 0.05). **F.** *Cross-species analysis of blood datasets*. Integration of human and mouse blood metabolomics data identifies shared dysregulated metabolites. Examples of commonly altered and significant metabolites include adenosine, 2-hydroxyglutarate, dodecanedioic acid, 2-hydroxycaproic acid, 3-ureidoisobutyrate, and L-threonic acid. **G.** *ASO treatment study in MDS mouse model*. MDS mice and WT littermates were treated with 350 µg of ASO targeting the human MECP2 copy. Samples were collected at weeks 4, 10, and 16 from the administration. A heatmap shows metabolite profiles that distinguish between untreated WT, untreated MDS, and ASO-treated MDS groups. **H.** *Shared and rescued metabolites*. Several metabolites dysregulated in both human and mouse MDS systemic dataset were responsive to ASO treatment in mice (“rescued” by ASO).

**Extended Data Fig. 3.** *Heat maps and multivariate analysis of neuro-metabolomics datasets***. A-B.** *Cerebrospinal fluid (CSF), brain organoids and cortex metabolomics*. Heat maps (**A**) showed distinct separation between MDS and control samples in CSF, brain organoids and cortex samples. **B.** Volcano plots of the corresponding datasets highlight significantly dysregulated metabolites. **C.** *Cortex metabolomics from ASO-treated MDS mice.* Mouse were treated with ASO targeting human copy of MECP2 and tissues collected after 4 weeks. Heat map and hierarchical clustering showed discriminating metabolites across conditions. **D.** *Pathway-level patterns in neurodatasets.* Metabolites associated with mitochondrial function (aconitic acid, succinate, 2 -Hydroxy-2-methylbutyrric acid) and nucleotide metabolism (inosine, uracil, hypoxanthine) exhibit similar trajectories in MDS across different models and tissues. **E.** *Cross-tissue integration of ASO response.* Intersection analysis of blood and neuro datasets identifies ASO-responsive metabolites common to both compartments, highlighting systemic metabolic correction following treatment. **F.** NADH and spermidine levels were elevated in MDS brain organoids, confirming altered redox balance, increased reductive pressure, and mitochondrial dysfunction in the disease model.

**Extended Data Fig. 4.** *Proteomic and transcriptome analysis performed in different models of MDS.* **A-B.** *GO enrichment of CSF proteome data.* **A.** Protein-level heat map with MECP2 motif annotation. Heat maps display the most significantly deregulated proteins in MDS CSF. Proteins in correspondence of green boxes contain predicted MECP2 binding motifs, suggesting direct transcriptional regulation. **B**. Gene Ontology (GO) analysis of CSF proteomic datasets revealed significant downregulation of ATP metabolic processes and nucleoside triphosphate metabolism in MDS samples (highlighted in red). **C.** Circular dot plots display specific proteins and metabolites associated with purine metabolism and TCA cycle pathways, identified through integrated analyses combining CSF metabolomics with proteomics, and brain organoid metabolomics with transcriptomics. **D**. Key enzymes of the tricarboxylic acid TCA cycle were significantly reduced in MDS compared to controls. These include ME1, ME3, SDHB, SUCLA1, and SUCLG1, indicating impaired TCA cycle in MDS. **E.** CSF proteome Volcano plot showed downregulated mitochondrial proteins (NDUFB10, GCSH) some involved in the electron transport chain (ETC), including BCS1L, and UQCRQ (highlighted in red). **F.** Transcriptome analysis of 60-day-old brain organoids revealed a downregulation of the glutathione detoxification pathway and an upregulation of the nucleotide biosynthesis pathway, as determined by Biocyc pathway analysis (*Enrichment analysis, Fisher exact test, p<0.05).* **G.** We next tested whether ASO treatment could restore metabolic balance. MSI (Spectroglyph, Inc.) shows increased succinic acid, a key TCA cycle product, in MDS compared to controls, which is normalized following ASO treatment (2.5 µM), supporting a direct link between MECP2 dosage and TCA cycle regulation. **H.** Mitomics analysis of 205 mitochondrial gene knockout (KO) lines revealed a greater number of upregulated TCA cycle metabolites compared to the downregulated ones, supporting an association between TCA metabolite accumulation and impaired mitochondrial function (T-test, *p<0.001*). For details, including specific genes deregulated in the TCA cycle and nucleotide metabolism pathways, see **Extended Data Tables 16–19**.

**Extended Data Fig. 5.** *Mitochondrial morphology and marker expression in MDS astrocytes, neurons, and brain organoids.* **A-B** Morphometric analysis of mitochondria in both astrocytes and neurons demonstrates a reduction in multiple structural parameters in MDS, including mean intensity, mean branch diameter, width, and Feret diameter (a measure of the longest axis of the mitochondrion), branch length along with diminished mean intensity (**A**). Additional parameters include mitochondrial perimeter which is significantly reduced in MDS neurons. In contrast, sphericity ratio is increased in MDS mitochondria, consistent with a shift toward more spherical mitochondrial shapes (**B**). **C.** 3D-rendering revealed altered mitochondrial morphology in MDS astrocytes. Quantitative 3D analysis showed a reduction in the number of mitochondria/cell, total mitochondrial surface area/cell, and mitochondrial branching/cell in MDS astrocytes compared to controls. *Scale bar= 10 μm.* For details see **Extended Data Video 1**. **D-E.** Immunostaining of mitochondrial markers COX4 (**D**), and SDHB (**E**) in 60-day-old brain organoids shows reduced expression in MDS samples compared to control ones, further supporting mitochondrial dysfunction in the MDS model. S*cale bar= 100 μm*. **F** Quantitative analysis of mitochondrial protein expression reveals significantly reduced mean intensity of COX4, and SDHB, in MDS organoids compared to control, consistent with reduced mitochondrial function. **G.** MSI (Tims-TOF Bruker, Inc.) revealed reduced ATP abundance in MDS organoids vs controls, supporting bioenergetic deficit in MDS brain organoids. *Scale bar= 5 mm*.

**Extended Data Fig. 6. A-D.** *Bioenergetic deficit in MDS.* Oxygen consumption rate (OCR) by Seahorse (**A**) and Resipher (**C**) platforms showed reduced level in MDS, demonstrates impaired mitochondrial respiration. Both basal and stimulated OCR levels are reduced, as well as ATP-linked respiration (**B**), calculated by subtracting post-oligomycin OCR from baseline, supporting defective ATP production. ***D.*** Expression studies of mitochondrial genes *ND1* and *ND2* is reduced in MDS organoids *(*p<0.05*), confirming its related mitochondrial mass. **E-H.** *Targeted Metabolite Supplementation as potential Driver of Mitochondrial Dysfunction.* 2-Hydroxyglutarate (2HG), a mitochondrial byoproduct of TCA, in controls organoids leads to a reduction of: *i.*mitochondrial mass (MitoTracker) (**E**), *ii.* membrane potential (TMRE) (**F**), *iii.* both basal and triggered oxygen consumption rate (**G**), *iv* as well as of ATP levels by Mass-spectrometry imaging (MSI-Tims-TOF Bruker, Inc.) (**H**), supporting a direct role of 2HG in mitochondrial dysfunction and mimicking the MDS phenotype. *Scale bar= 4 mm*. **I-K**. *CIII impairment validation using CUT&RUN datasets.* CUT&RUN experimental data demonstrated that several proteins associated with Complex III (CIII) show direct MECP2 binding, supporting both biological findings and bioinformatic predictions, including UQCRFS1 and UQCRB (**I**). Volcano plot depicts peaks significantly upregulated in the MDS mouse model compared to controls (**J**). MECP2 binding analysis by CUT&RUN reveals that numerous CIII-related genes are significantly bound by MECP2 in MDS samples, whereas no MECP2 binding is detected in MECP2 knockout (KO) models, suggesting a direct regulatory role of MECP2 in mitochondrial gene expression (**K**). **L.** NAD⁺, FAD, and oxidized CoQ10 levels were reduced in MDS brain organoids, as measured by mass spectrometry imaging (MSI; Spectroglyph, Inc.), consistent with metabolic alterations expected under CIII deficiency conditions.

**Extended Data Fig. 7. A.** *Electron Transport Chain Impairment links Mitochondrial Deficits to MDS.* Comparative protein profiling of CSF and brain organoids reveals global reductions in ETC-related proteins in MDS. In addition to the consistent downregulation of CIII in both CSF and organoids proteome, shared reductions are observed in complex IV (CIV) and ATP synthase (complex V, CV), highlighting widespread disruption of mitochondrial oxidative phosphorylation. **B-C.** A specific reduction of PPOX levels in CSF (**B)** and brain organoids (**C**) further supports CIII deficiency, as PPOX activity depends on mitochondrial electron transport through CIII for proper function. **D-G***. Oxidative stress is increased in MDS brain organoids.* **D.** Induced glutathione (GSH) response following H2O2 pulse is significantly blunted in MDS organoids, indicating impaired antioxidant capacity. **E**. Pathway-enriched analysis showed that glutathione metabolism was reduced in both radial glia and neuronal clusters of MDS vs controls (Wilcoxon rank-sum test, *\*p<0.05,* ****p<0.001). **F.** Volcano plots of RNA-seq at 30 and 60 days. Volcano plots show consistent upregulation of *MECP2* and *IRAK1*, genes located in minimal overlapping regions associated with MDS pathology, along with *XRRA1.* **G.** CellRox fluorescent mean intensity, a marker of oxidative stress, measured by flow cytometry, is elevated in MDS (*T-Test, ****p<0.001)*. **H-J.** γH2AX foci, indicative of DNA damage, are significantly increased in MDS organoids compared to controls, as shown by quantification of γH2AX foci over nuclei, total number of Edu^+^ nuclei, (**H**) and representative low-(**I**, *scale bar = 50 μm*) and high-resolution (**J**, 100×, *scale bar = 10 μm*) immunofluorescence images.

**Extended Data Fig. 8. A.** Pathway enrichment analysis demonstrates that the DNA damage checkpoint pathway is significantly upregulated in both neuronal and radial glia clusters in MDS organoids (Wilcoxon rank-sum test,*\*\*\*\*p<0.001)*. **B-C.** Single-cell RNA-seq shows upregulation of DNA-damage genes (*CCND1*, *BRCA1*; B) and DNA damage response (DDR) markers (*TP53*, *ATR*) in radial glia, with *TP53* also elevated and *ATR* trending upward in neuronal clusters (violin plots **C**). **D-E.** CRISPR *CYC1 KO* (a component of mitochondrial Complex III) in control lines leads to elevated levels of both TCA and purine metabolites (**D**) along with increased protein levels of ADSL, PFAS, and IMPDH2 (**E**), further linking mitochondrial dysfunction to activation of purine biosynthesis. **F-G.** Gene expression analysis confirms that multiple genes involved in purine metabolism are significantly upregulated in MDS. Transcriptome revealed increased level of genes associated to de novo pathway (*ADSS2, ENPP2, ENPP3*), while reduced levels of gene associated to salvage pathway (*SLC29A2*) *(T-Test *p<0.05, **p<0.01*) (**F**). Single-cell RNA-seq further showed activation of the pentose phosphate pathway (**G**), which provides essential precursors and reducing power required for activation of the purine biosynthetic pathway.

**Extended Data Figure 9.** *Further evidence of purine pathway upregulation in MDS organoids.* **A-B.** Immunofluorescence staining for PFAS (**A**) and corresponding quantification (**B**) reveal enhanced expression of this key enzyme in the de novo purine synthesis pathway, co-stained with the mitochondrial marker TOMM20. PFAS intensity is increased in both SOX2⁺ neuroprogenitor and SOX2⁻ neuronal regions of MDS organoids, while TOMM20 signal is reduced, suggesting a shift from oxidative metabolism toward nucleotide biosynthesis under metabolic stress. Together, these results further corroborate purine pathway upregulation in MDS. *Scale bar = 100 µm.* **C-D.** Western blot analysis demonstrates significantly elevated PFAS protein levels in MDS organoids, consistent with transcriptomic findings. **E-G**. Quantification of purine biosynthesis enzymes revealed increased GART expression (E; corresponding to Main Figure 5A) and elevated PFAS area and mean intensity per purinosome (F; corresponding to Main Figure 5B) in MDS organoids, indicating activation of the de novo purine biosynthesis pathway. PFAS levels were further assessed following ASO treatment (G; corresponding to Main Figure 5C), showing restoration toward control levels, consistent with recovery of metabolic balance. **H**. Pseudotime trajectory analysis shows increased expression of *ADSL, NT5C1B, ADSS2, GART, AMPD1*, and *GUK1* across multiple developmental time points in MDS organoids. This coordinated upregulation indicates activation of the de novo purine biosynthesis pathway, linking TCA dysfunction to enhanced nucleotide synthesis under *MECP2* overexpression. **I.** Specific purinergic receptors and pathway determinants, including ADORA, P2RX3, SLC25A13, and PDE4B, were increased in MDS organoids based on single-cell RNA-seq analysis, supporting upregulation of the purinergic signaling pathway.

**Extended Data Figure 10. A-B**. EdU labeling after one week of treatment reveals impaired neuroprogenitor migration within rosettes in MDS organoids compared to controls (**A**), as evidenced by a reduced number of DCX⁺/EdU⁺ double-positive cells (**B**), indicating defective neuroblast generation and migration from the apical progenitor zones. *Scale bar = 50 µm.* **C-F.** *Disrupted Neuroprogenitor Niche Dynamics in MDS.* **C.** Quantification of rosette area from Figure 6A shows a reduction in MDS, consistent with diminished proliferation and/or aberrant differentiation of neural progenitors. **D.** Single-cell RNA-seq trajectory analysis demonstrated reduced neuronal maturation in MDS organoids compared to controls, consistent with impaired neurodevelopment. **E-F.** Neurometric analysis revealed a trend toward increased total neurite length per neuron (**E**) and a significant increase in average branches per neuron (**F**), supporting enhanced neuronal branching in MDS organoids, consistent with hyperactivation of the purinergic signaling pathway. **G.** Immunostaining for phosphorylated vimentin (p-vimentin), a marker of dividing radial glial cells, showed a reduced number of positive cells in MDS organoids, further supporting defects in progenitor proliferation and maintenance. *Scale bar= 25 μm.* **H.** Expression of proliferation markers (*TOP2A, MKI67*) and neural progenitor marker (*PAX6*) was significantly downregulated in MDS radial glia cluster, indicating impaired progenitor maintenance and cell cycle progression (Wilcoxon rank-sum test, *\*\*p<0.01,* ****p<0.001). **I-J.** *Linking metabolomic alterations to cell fate changes in MDS brain organoids.* **I.** MSI (Spectroglyph, Inc) revealed significantly higher levels of glutamate and glutamine in the rosette regions of controls compared to MDS. Both metabolites are tightly linked to the TCA cycle. *Scale bar= 1 mm.* **J.** Consistently, the glutamatergic pathway was downregulated in the MDS radial glia cluster, indicating that TCA cycle–linked metabolic dysregulation and associated damage manifest early in progenitor development (Wilcoxon rank-sum test, *\*\*p<0.01,* ****p<0.001).

## METHODS

### Participants and samples obtained

#### Participants

The study was approved by the Institutional Review Board at Baylor College of Medicine (H-46532) and was performed in accordance with the World Medical Association Declaration of Helsinki. We recruited 30 subjects with MDS syndrome confirmed by the Chromosomal Microarray (29 males,1 female; average age 10.48 years, range 1.9-31 years) and 32 healthy controls (26 males, 6 females; average age 39.19 years, range −66 years) who were family members (mother, fathers and siblings) to minimize genetic and environmental confounders. In addition, we recruited 5 typically developing individuals who had a clinical indication for a spinal tap (Traumatic brain injury, Idiopathic intracranial hypertension, Aqueductal stenosis related hydrocephalus). All MDS participants had comprehensive clinical, neurological, developmental, and genetic assessments. Before sampling, a 48hr dietary intake was obtained, including intake of all medications and supplements. Please see **Extended Data Fig. 1A** and **Tables 1-3** for details.

#### Sample collection

i. *Blood* samples were collected for all MDS participants (10 had blood draw twice, 6-8 months apart; total number of samples analyzed n=40) and their family controls (n=32) after overnight fasting and before morning medications (**Extended Data Table 1,3**). Blood was collected in Eppendorf tubes with 20% deuterated EDTA. Samples were centrifuged at 1800g for 10min at 4°C to separate plasma, followed by second centrifugation at 300g for 15min at 4°C, to discard cell debris. Samples were aliquoted at 300 μl and stored at −80°C
ii. *CSF* was collected from 8 MDS participants twice, 6-8 months apart (total number of samples n=16) and 5 typically developing individuals (one sample run twice), under sedation and in presence of the Texas Children’s Hospital Anesthesia team. Following spinal tap, CSF was immediately aliquoted and stored at −80°C.
iii. *Skin biopsy* was obtained from four male MDS participants who had similar extent of duplication ranging from 232-1200 kb (**Extended Data Tables 1, 10)**. Skin biopsies were processed into fibroblasts by the IDDRC Tissue Culture Facility Core at Baylor College of Medicine. Briefly, they were rinsed in DPBS and placed at the bottom of a 6 cm tissue culture dish. Biopsies were minced between two sterile scalpel blades. Alpha-MEM +10% fetal bovine serum (FBS) + 1× Penicillin–Streptomycin was placed on top, and minced material was cultured at 37°C in 5% CO_2_ and 5% O_2_ incubator. Cultures were monitored for fibroblast cell outgrowth. When the cells sufficiently expanded, the culture was trypsinized and expanded in T25 and T75 flasks until ready for further processing.

### Animal models

In this study, we used the transgenic *Mecp2*-overexpressing mouse model^103^ (MDS mice, harboring one copy of human *MECP2* in addition to the endogenous mouse gene), their wild-type (WT) littermate controls. All experiments were performed in accordance with the eighth edition Guide for the Care and Use of Laboratory Animals and approved by the Ionis Pharmaceuticals Institutional Animal Care and Use Committee (IACUC). The mouse studies were done according to the IACUC protocol 2021-1176. These rodent model were treated by antisense oligonucleotides targeting *MECP2* (see below).

#### ASO administration in mice

18, 8-12-week-old, male mice were treated with 350 µg ASO targeting human *MECP2*. The scalp was shaved and the mouse was positioned on a stereotaxic instrument under anesthesia at 2% isoflurane. The scalp was scrubbed with betadine and cleaned with ethanol, and a 1-2 cm incision was made along the mid-sagittal line to expose the skull. The subcutaneous and periosteal membranes were scraped away from the skull with sterile cotton swabs. A sterile Hamilton gas tight microsyringe with an attached 22-26 gauge needle held by a stereotaxic micromanipulator was used to inject the ASO into the ventricles at an approximate rate of 1 µL/second. The tip of the needle was passed through the skull 0.3 mm anterior, 1 mm lateral, and 3 mm ventral in reference to bregma. The needle was withdrawn after an approximately 3 min and the skin was sutured with 5-0 nylon suture. The mouse was removed from the stereotaxic apparatus to its home cage to recover from anesthesia.

#### Blood collection

Whole blood was collected via cheek bleed (submandibular) and placed into a serum tube (Greiner Bio-One catalog number 450470). Blood was allowed to clot for 30 min and was spun for 10 min at 1500g thereafter. Serum was collected and stored at −80°C until analysis. Serum from 48 WT, 35 MDS mice, 18 MDS mice treated with ASO was collected after 4,10 and 16 weeks.

#### Tissue collection

Mice were deeply anesthetized using CO_2_ and then euthanized by rapid decapitation. The brain was dissected, and fine forceps was used to dissect out the cortex, which was snap frozen and stored at −80°C until analysis. Cortices were collected from 6 WT mice, 8 MDS mice, and 6 MDS mice treated with ASO.

### iPSCs and organoid models

#### iPSC generation

iPSC were generated from fibroblasts of 3 male patients and GM23815, GM01888, and GM04545 male control lines, by Sendai virus reprogramming^102^ (**Extended Data Table 10)**. Their pluripotent status was confirmed by alkaline phosphatase staining and pluripotency score (ThermoFisher), while absence of chromosomal aberration was checked by Karyostat (ThermoFisher) (**Extended Data Figure 1B**). At least 3 clones per each line were banked and each clone used underwent pluripotency confirmation and Karyostat (Thermo Fisher) every 10 passages. All lines tested negative for Mycoplasma every 10 passages.

#### Cortical organoid generation

We generated cortical organoids from iPSC lines (n=3 male controls, n=4 male MDS patients) using a modified Pasca protocol^104^. Briefly, iPSC colonies were dissociated using Accutase (Thermo Fisher Scientific) for 10 min at 37°C. After centrifugation for 3 min at 270*g*, the dissociated cells were resuspended in Basal media supplemented with 10 µM SB431542 (Stemgent), 1 µM dorsomorphin (R&D Systems) and 5 µM Rho kinase inhibitor (Y-27632; Calbiochem, Sigma-Aldrich). 1.5 million cells were transferred into low-attachment AggreWell 800, pre-treated with Anti-Adherence Rinsing Solution. To achieve rapid and efficient neural induction, both the BMP and TGF-b signaling pathways were inhibited with small molecules dorsomorphin and SB-431542 for the first 5 days of the culture. After 8 days in the Aggrewell plate, the floating spheroids were collected into suspension dishes and Neurobasal/DMEM/F12 with B-27 without Vit A and N2 was supplemented with FGF2 and ascorbic acid up to day 21. To promote differentiation of neuroprogenitors into neurons, brain-derived neurotrophic factor (BDNF) and neurotrophic factor 3 (NT3) were added starting on day 22 until day 60. From day 60 onward, only neural medium without growth factors was used for media changes every other day. Organoids were collected for downstream analysis at 30 and 60 days. Comparative analysis across time points (n = 6 samples) revealed that the greatest differences between MDS and control organoids were observed at 60 days. Therefore, the 60-day time point was selected for all subsequent downstream analysis.

#### Organoid disaggregation

Organoids were disaggregated into individual cells using the Neural Tissue Dissociation Kit (Miltenyi Biotec, NRW, Germany). Cells were washed twice, counted, and assessed for viability using trypan blue exclusion, then either replated for staining, processed for flow cytometry, or prepared for single-cell RNA sequencing.

### Antisense oligonucleotides

ASOs were synthesized by Ionis Pharmaceuticals. The ASOs consisted of 20 chemically modified nucleotides [2′-*O*-(2-methoxyethyl) (MOE) gapmer] with a central gap region of 10 deoxynucleotides that was flanked on both its 5′ and 3′ sides by five MOE-modified nucleotides. The ASO backbone included phosphodiester (PO) and phosphorothioate (PS). Reconstituted ASO was stored at 100mg/mL stock in DPBS −/− then diluted to desired dosing concentration in the same DPBS. Reconstituted ASOs were stored at −20°C. Both ASOs are 5-10-5 MOE MBB (EEoEoEoEo-10-EoEoEEE where E is MOE and O is phosphodiester). A dose of 350 µg was used and mecp2 expression reduction was confirmed in all models (**Extended Data Fig. 1).**

We next tested if treatment with an ASO that targets *MeCP2* for RNase H1-mediated degradation (*MECP2* ASO) was effective also in MDS human brain organoids. Ionis Pharmaceuticals designed and synthesized the *MECP2* and control ASOs^105^. From 5′ to 3′ the linkages are 1-PS, 4-PO, 10-PS, 2-PO, and 2-PS. ASOs were added directly to the media to achieve a final concentration of 2.5 µM, administered in two doses every other day. Organoids were collected two weeks after the first treatment (**Extended Data Fig 7C**). Both control and MDS organoids were treated with ASOs, with blank media (no supplements) used as a vehicle control.

*ASO for MDS mouse:* TTCATTTCTCTTGTTTCGCA.

*ASO for cortical organoids:* GTTCAATATGTCATCCGAAG^16^

Control (non-targeting) ASO: CCTATAGGACTATCCAGGAA

### Metabolomics

#### LC-MS/MS sample preparation

20 µl of *plasma/serum* or 40 µl of *CSF* was mixed with 100 µl or 80 µl of ice-cold methanol respectively, and 0.5 µM internal standard composed of L-DCA-d5 (negative mode) and Agomelatine (positive mode). The samples were vortexed twice and spun down at 15,000 rcf for 20 min at +4°C. After centrifugation, supernatant (40-80 µl) was removed and transferred into LC vials, vortexed again, and used for analysis. For quality control (QC) samples, 3 μl from each sample was pooled and mixed with 5x volume of ice-cold methanol, then processed using the same protocol as individual samples.

*Mouse cortices* from WT and MDS (treated and untreated) were dissected and homogenized, and the resulting lysates were processed for metabolite extraction. Samples were treated with 50% methanol, homogenized, and centrifuged, after which the supernatants were collected and analyzed.

*Brain organoids* were smashed in PBS, sonicated to get lysates, and protein content was measured using Pierce BCA Protein Assay Kit. 300 µL of organoid lysate was normalized at concentration of 1 µg/µl (total protein 300 µg) and dried in a vacuum concentrator at room temperature. The samples were reconstituted in 24 µL of 50% methanol (containing 0.05 µM of agomelatine and 0.25 µM of lithocholic acid-d5 as the positive and negative internal standards, respectively). The samples were vortex for 1 min, centrifuged at 15000 rcf for 15 min at 4°C, and 20 µL of the supernatant was transferred to the sample vials for LC-MS analysis. The sample protein concentration post concentration is 12.5 ug/uL.

Samples were analyzed using UHPLC coupled with Q Exactive MS (Thermo Fisher Scientific, San Jose, CA) equipped with a 100 mm x 2.1 mm BEH C-18 column (^106^)). The column temperature was set at 40°C and the 0.3 ml/min of flow rate was used with a gradient ranging from 2% to 98% aqueous acetonitrile containing 0.1% formic acid in a 15-min run. Q Exactive MS was operated in full scan and DDA-MS/MS with electrospray ionization in positive and negative modes. MS data were acquired from 80 to 1200 m/z with the resolution set to 140,000, the AGC target at 106, and the maximum injection time at 100 msec in profile mode. QC samples were used multiple times during sample runs.

#### Data analysis

The primary data were processed and normalized by Compound Discoverer 3.2 (Thermo Fisher Scientific, San Jose, CA)^107^ and SIMCAP14 (Umetrics, Kinnelon, NJ) software to generate a multivariate data matrix. Manually curated peaks were annotated based on in-house database of m/z and retention time of external standards; only peaks that passed the experimental ion ratios for the product ions (where more than one product existed) were integrated and peak area values were then exported. Compound Discoverer performs retention time alignment, QC normalization, unknown compound detection, and compound grouping across all samples predicts elemental compositions for all compounds, fills gaps across all samples, and hides chemical background. Multivariate classification models were built using orthogonal projection to latent structures-discriminant analysis (OPLS-DA), conducted on Pareto-scaled data for determining the primary metabolites contributing to group differences. Compound Discoverer software streamlines compound identification and comparative analyses and provides extensive filtering and data visualization capabilities to process untargeted metabolomics data by the removal of adducts, fragments, naturally occurring isotopes, and other confounders.

Differential analysis with p <0.05 and log2 fold change of 0.6 was considered significant. Comparative list of identified metabolites from all datasets is included in **Extended Data**: systemic dataset from human (**Extended Data Table 5**) and mice (**Extended Data Tables7**), human CSF (**Extended Data Table 11**), brain organoids (**Extended Data Table 12**) and mouse cortex (**Extended Data Tables 13**).

MS/MS analysis was run for organoids samples. The exact mass and predicted formula and structure of the metabolite obtained from Q Exactive MS was used to query metabolomics databases (e.g., HMDB, Metlin). To map identified and cross-referenced compounds to biological pathways, we utilized established metabolomic databases, including Kyoto Encyclopedia of Genes and Genomes (KEGG) and Human Metabolome Database (HMDB). For the conversion of compound names to KEGG identifiers, the MetaboAnalystR v6.0^108^ ID conversion tool was used (https://www.metaboanalyst.ca/MetaboAnalyst/upload/ConvertView.xhtml).

#### Batch correction

LC–MS peaks, characterized by their mass-to-charge ratio (m/z) and retention time (RT), are highly sensitive to experimental conditions and analytical batch effects. Since plasma and CSF samples were analyzed in different batches, we applied the ComBat method for batch correction, aiming to minimize technical noise while preserving true biological differences.

#### Age as co-variable

To assess the potential influence of age, each molecular feature was modeled as a function of chronological age using linear regression. The coefficient of determination (R²) quantified the fraction of variance explained by age, and false discovery rate (FDR) correction was applied to identify significant age-associated features. However, when groups differ completely in age with no overlap, age and diagnosis become perfectly collinear. In such cases, age correction is not valid because any apparent difference may reflect developmental rather than disease-specific effects. For these reasons, age effects were evaluated separately within each group. In controls (all adults), no significant associations were detected between age and metabolite levels, indicating stable metabolic profiles across adulthood. Among significant metabolites in MDS organoids three showed modest negative correlations with age; however, given the lack of age overlap with controls, these trends most likely reflect disease-stage–dependent metabolic shifts rather than true age effects. The convergence of findings across multiple systems—including targeted and untargeted plasma metabolomics, cerebrospinal fluid, cortex, and patient-derived organoids—together with parallel results in the MDS mouse model and ASO dependency, indicates that these metabolic alterations are independent of age. The absence of age-related effects in controls, combined with the consistent directionality of metabolic changes across species, platforms, and sample types, further argues against age as a confounding variable. Although a partial adaptive remodeling of the MDS metabolome over time cannot be excluded, this effect is likely modest, as familial controls were selected to minimize dietary and environmental variability.

#### One factor analysis

Statistical significance was assessed using two-tailed Student’s *t*-tests to evaluate whether each metabolite was significantly associated with the outcome of interest, unless otherwise noted. For comparisons involving two or more groups, one-way ANOVA was used to assess group differences (Prism 9.5.1, Boston, MA, USA). Analyses yielding *p* < 0.05 were considered statistically significant. Metabolite intensity values were log2-transformed prior to visualization to improve normalization and reduce skewness, which is common in omics data. This transformation also helps stabilize variance across samples. All the plots display the mean and standard error of the mean (SEM) of log2-transformed intensities for each condition, highlighting both central tendency and variability. To discriminate between groups, we applied both univariate and multivariate statistical approaches. Univariate analyses included volcano plots and correlation analyses. Multivariate analyses comprised Principal Component Analysis (PCA), Partial Least Squares–Discriminant Analysis (PLS-DA), and hierarchical clustering (heatmaps) (Top 50 metabolites, euclidean measure, clustering method average). Important features were selected using one-way ANOVA and post-hoc analysis with a *p*-value threshold of 0.05. For hierarchical clustering, we used Euclidean distance as the metric and Ward’s linkage as the clustering algorithm, which minimizes the total within-cluster variance. In PLS-DA, we calculated Variable Importance in Projection (VIP) scores, representing a weighted sum of squares of the PLS loadings, weighted by the proportion of explained Y-variation in each component.

#### Metadata analysis

To improve the identification of robust and reproducible metabolic biomarkers, we performed horizontal integration—commonly referred to as metabolomic meta-analysis—across multiple independent studies investigating the same condition in comparable populations. As a preliminary step, differential expression analysis was performed independently for each dataset using linear modeling with the Limma package, allowing for the identification of candidate metabolites and the evaluation of study-specific trends. Statistical meta-analysis integrated human and mouse plasma with brain-related samples (human CSF, mouse cortex, brain organoids), retaining only consistently dysregulated (p-corrected) metabolites across datasets. Meta-analysis was then performed using a p-value combination approach based on Stouffer’s method, which integrates both effect direction and magnitude while weighting by study size, under the assumption of comparable data quality. For blood datasets (human plasma and mouse serum), we applied a significance threshold of *p* < 0.05; for neuro datasets (human CSF, brain organoids, and mouse cortex), a more permissive threshold of *p* < 0.25 was used to accommodate (i) the inclusion of three distinct metabolomic datasets and (ii) the extracellular nature of CSF, where metabolite levels may display trends opposite to those in intracellular compartments (mouse cortex and human brain organoids) due to compartmental shifts or differences in extraction efficiency. Meta-analysis outputs were then cross-referenced with metabolite changes observed after ASO treatment. Metabolites showing significant alteration in the opposite direction following ASO treatment—prioritized when correlation exceeded 0.25 according to *pattern hunter* predefined trends—were considered potential rescue targets. Please see **Extended Data Tables 9,13** for details.

#### Enrichment analysis

To identify biologically relevant pathways in MDS, we performed enrichment analyses on metabolites from the meta-analysis using MetaboAnalystR v6.0 (https://www.metaboanalyst.ca/MetaboAnalyst/ModuleView.xhtml)^109^ and Human BioCyc (https://biocyc.org/). Metabolite Set Enrichment Analysis (MSEA) detected coordinated alterations across predefined metabolite sets, capturing subtle, consistent shifts missed by threshold-based approaches. Over-Representation Analysis (ORA) with the hypergeometric test determined pathway enrichment, with one-tailed p-values adjusted for multiple testing. Enrichment scores were calculated on ASO-filtered blood datasets, while CSF, brain organoid, and cortex datasets were analyzed independently before identifying shared pathways. In detail, enrichment analysis models to compute the theoretical number of hits by random chance (expected hits), and compare this number with what we actually observed based on the data (observed hits) (*q*/Expected *q*) (**Figure 1I, J**). Integrating blood and neuro datasets revealed convergent metabolic dysregulation in tricarboxylic acid and nucleotide metabolism (**Extended Data Tables 14-15**).

#### Multiomics data integration and Joint Pathway Analysis

— To overcome limitations in metabolomic data interpretation and derive biologically meaningful insights, we integrated metabolite profiles with transcriptomic and proteomic datasets from brain organoids and CSF, respectively, at the pathway level. Enrichment analyses were performed using the hypergeometric test, and data integration was achieved through degree centrality and combined-query methods. This multi-omics framework provided a comprehensive overview of molecular alterations in MDS, enabling a more robust and biologically coherent interpretation of complex datasets. Using MetaboAnalyst 6.0^109–111^ we projected metabolites and genes onto biological networks, and conducted joint pathway analysis using the Interaction Network Explorer, generating a global view of interconnected metabolic and signaling pathways. This analysis combined enrichment and topology-based methods using the KEGG metabolic pathway database, integrating univariate statistics, over-representation analysis, and advanced algorithms (Global Test, GlobalAncova, and network topology metrics). We applied this framework to:

(1) CSF metabolite data integrated with proteomics, and

(2) brain organoid metabolomes integrated with transcriptomic profiles (see **Extended Data Tables 16–17**). Specifically, we used the hypergeometric test with centrality degree and combined query functions for enhanced sensitivity. Pathway validation using BioCyc perturbation scores ranked pathways by Pathway Perturbation Score (PPS) and Differential Pathway Perturbation Score (DPPS) to assess overall regulation. Joint pathway analysis revealed convergent disruptions in nucleotide and mitochondrial pathways across both CSF– proteomics and organoid–transcriptomics datasets, findings that were corroborated by BioCyc-based statistical tests (Fisher’s exact test, *p* < 0.1; see **Extended Data Tables 18-19** for details).

### Biomarkers validation: Plasma samples preparation

#### LC-MS

20 µl of plasma samples was mixed with 100 µl of methanol and 0.5 µM internal standard composed of L-DCA-d5 (negative mode) and Agomelatine (positive mode). We then vortexed the sample and spin it at 15000 rcf for 15 min at +4 C. After centrifugation, supernatant (40-60 µl) was removed and transferred into LC vials, vortexed again, and put into the UHPLC/Orbitrap MS for analysis. For quality control, we took at least 3 µl from each plasma sample supernatant and mixed all samples together (40-60 µl) for analysis. Prior to sample analysis, blanks injections were performed to calibrate the instrument to ensure that there are no impurity signals or contamination. Then, the QC samples were run to determine the signal range coming from all plasma samples together. Then the individual plasma samples were run. Ensuring that the instrument is stable prior to sample analysis helps maintain consistency between the runs.

We used a Thermo Q Exactive Orbitrap mass spectrometer coupled to Vanquish Horizon ultra-high performance lipid chromatography (UHPLC) system. Q Exactive MS was operated in full scan with electrospray ionization in positive and negative modes. MS data were acquired from 80 to 1200 m/z with the resolution set to 140,000 and 5 ppm mass accuracy, the AGC target at 10^6^, and the maximum injection time at 100 msec in profile mode. These settings have been used to acquire high resolution MS1 data on human plasma samples. In addition, blank samples and pooled reference sample (QC) were analyzed in MS1 mode. The exact mass and predicted formula and structure of the metabolite obtained from Q Exactive MS was used to query metabolomics databases (e.g., METLIN, HMDB) or chemical databases (e.g., Chemsipder). Some biomarkers that distinguished MDS from controls using both human and mice plasma, and that were corrected by ASO in MDS mice to control levels were some mitochondria related pathway (2HG at negative mode) and nucleotide related metabolites (adenine, adenosine monophosphate, at positive ion mode, upregulated in respectively in human and mice plasma and brain-related samples). We validated biomarkers in positive mode using HILIC column, and biomarkers in negative mode using BEH C-18 column. We validated all biomarkers listed above. The standards of the 3 identified biomarkers were dissolved in methanol, mixed and diluted to 5 ug/mL in 50% methanol. The mixture of standards (3 uL) was injected before the samples to confirm the retention times of the analytes. We then ran blanks to check for impurities, then the mixture of standards, and then 3 uL of individual samples. The analytes were separated on a Phenomenex Kinetex HILIC column (1.7 u, 100*2.1mm) for positively ionized standards and a 100 mm x 2.1 mm BEH C-18 column for negatively ionized standards, analyzed by the Q Exploris 120 Orbitrap MS coupled with a Vanquish UHPLC. The mobile phases consisted of water (A, containing 0.1% formic acid) and 90% acetonitrile (B, containing 10 mM of ammonium formate). The flow rate was set at 0.4 mL/min and the gradient was from 99% B to 40% B in an 18-min run for HILIC column The C-18 column temperature was set at 40°C and the 0.3 ml/min of flow rate was used with a gradient ranging from 2% to 98% aqueous acetonitrile containing 0.1% formic acid in a 15-min run for negative mode. Q Exploris was run under the positive and negative full-scan profile mode with the mass range of m/z = 70-500. The MS resolution was set at 120,000 and RF lens 50%. The spray voltage was set at 3500 V. The capillary and vaporizer temperatures were set at 325 and 350 degrees, respectively. Ultra-pure nitrogen was used as the sheath (50 arbitrary units), aux (10 arbitrary units) and sweep (1.0 arbitrary units) gases. The peaks of the analytes were extracted with their exact masses (mass error 5 ppm) and were integrated for relative quantification (**Extended Data Figure 2B**).

#### Targeted metabolomic profiling (GMAPs)

Plasma samples were analyzed using the Global Metabolomics-Assay Platform (GMAPs), a targeted mass spectrometry–based method previously described^112^. Quantitative z-score profiling revealed increased levels of malate (z = 0.25), 2-hydroxyglutarate (z = 1.29), 4-hydroxy-2-oxoglutaric acid (z = 0.88), inosine monophosphate (z = 0.45), adenine (z = 0.79), and succinyladenosine (z = 0.56). The abundance pattern of these metabolites aligned with the alterations identified in the untargeted metabolomic analysis, reinforcing the robustness of the metabolic signature in MDS plasma (**Extended Data Table 6**).

### Mass spectrometry imaging (MSI)

For MSI, we used both Spectroglyph (Spectroglyph LLC, United States) and Tims-TOF (Bruker, Inc.; for higher resolution imaging) approaches.

#### MALDI MSI

A first subset of experiments was run using a high resolution MSI platform, by mounting a MALDI ion source containing a dual-ion funnel interface (Spectroglyph LLC, United States) to a Q-Exactive mass spectrometer (Thermo Fisher Scientific, Massachusetts, United States) as previously described^36^. The Orbitrap mass spectrometry was operated at ion injection time of 250 ms and Fourier Transform MS (FTMS) spectra were acquired in a profile mode using a target mass resolution of 70,000 (Full Width Half Maximum (FWHM) at m/z 400, m/z = mass-to-charge ratio). During MALDI imaging, mass spectrometer starts to acquire data after switching on to contact closure. The signal is sent from the MALDI injector and communicated via the ‘Peripheral Control’ input connection at the side of the Q Exactive MS, and the automatic gain control was switched off.

#### MALDI-TOF MSI

For a second set of experiments a high resolution MSI platform for visualizing small metabolites is implemented by a timsTOF fleX MALDI-2 mass spectrometer (Bruker Daltonics, Bremen, Germany). MALDI-MSI spectra were acquired in the negative ion mode from 80–900 Da. Transfer settings were 250 V peak-to-peak (Vpp; funnel 1 RF), 200 Vpp (funnel 2 RF), and 200 Vpp (multipole RF). Focus pretime-of-flight (TOF) transfer time was set at 80 μs and prepulse storage at 5 μs. The quadrupole ion energy was 5 eV with a low mass of m/z 50. Collision cell energy was 10 eV with the collision RF set to 800 Vpp. All the spectra were recorded using a 0.5 kHz laser repetition rate with 600 laser shots accumulated at each pixel with MALDI-2 off. The smart beam was set to a single focused beam at 40% power with a scan range of 20 μm ×20 μm. External calibration was completed using an Agilent Tuning Mix for ESI-TOF.

#### Tissue preparation

The tissue preparation was the same for both MSI approaches. Brain organoids were embedded in 10% cold fish skin gelatin, gently frozen, and sectioned at 14 µm, as previously described^36^.

#### Matrix preparation and application

Immediately after taking the slides with tissue sections out of the freezer, tissue sections were put in a desiccator for 20 min to minimize condensation of atmospheric water on their surfaces. After desiccation, we used *N*-(1-naphthyl) ethylenediamine dihydrochloride (NEDC)^36^, 10 mg/ml in 70% methanol, to spray and coat sections with heated pneumatic sprayer (HTX Technologies LLC, Carrboro) (**Extended Data Table 22)**.

#### Data analysis

In both MSI approaches the images were analyzed by SCiLS Lab MVS Version 2023a Core (Bruker Daltonics, Germany) employing total ion count (TIC) normalization. The Control and MDS MSI data files were differentiated by spatial identity of small metabolites (TCA, Glycolysis and purines) by which regions of organoids were mapped. The raw data file was uploaded to metabolite identification of the discriminated m/z values by employing the Human Metabolome Database (HMDB) (tolerance <5 ppm). Metabolic pathways were assigned based on identified m/z values.

### Transcriptomics

#### Bulk RNA sequencing (RNAseq)

RNA was isolated as described above and sent to Genomic and RNA Profiling Core (GARP) at Baylor College of Medicine for RNA integrity assessment, library preparation, and sequencing on the Illumina HiSeq platform. For each sample, approximately 100 million 150 bp pair-end reads were generated. Raw reads were trimmed before mapping by Trimmomatic v0.39 using the adapter reference TruSeq3-PE.fa:2:30:10.

Trimmed reads were aligned to GRCh38.p12 version 28 human genome assembly from. Read counts were then normalized and analyzed for differential gene expression using the DESeq2 package v1.34.0. Two initial baseline comparisons were made, each normalized independently. Genes were considered a commonly differentially expressed gene if the p-value was <0.05.

#### Bulk RNA-seq data analysis

We quantified transcript-level abundance using Salmon (v1.2.0) with default parameters and the GRCh38 reference genome (Ensembl v108). Following a thorough quality assessment, one control sample was excluded from downstream analysis due to data quality concerns. To ensure robust statistical analysis, we retained genes with a minimum read count of 10 across all samples prior to performing differential expression analysis. The differentially expressed gene (DEG) analysis was performed using the R package DESeq2 (v1.38.3). DEGs between the MDS and control groups were selected based on an absolute log2 fold change > 0.58 and an adjusted p <0.05. The pathway analysis was performed using the R package clusterProfiler (v4.6.2) with the function enrichGO (OrgDb = org.Hs.eg.db (v3.16.0), pAdjustMethod = “BH”, pvalueCutoff = 0.05, qvalueCutoff = 0.05) and the function enrichKEGG (organism = ‘hsa’, pAdjustMethod = “BH”, pvalueCutoff = 0.05, qvalueCutoff = 0.05). The up-regulated and down-regulated DEGs were used as input to identify up-regulated and down-regulated pathways (GEO repository: GSE304672).

#### Single Cell RNAseq (scRNAseq)

The disaggegated organoids were fixed according to the manufacturer’s protocol (Parse Biosciences), including 0.5% BSA in the fixation solution. After fixation, the cell concentration was measured for each suspension before storing at –80°C. For barcoding, equal numbers of cells from each suspension were used with the Evercode Whole Transcriptome kit (Parse Biosciences) following manufacturer’s instructions. Barcoding and sequencing library preparation were performed according to the protocol, resulting in six sub-libraries. The quality and fragment size distribution of the final sub-libraries were assessed using a Bioanalyzer (Agilent Technologies, Santa Clara, CA) and quantified with a Nanodrop spectrophotometer. The sub-libraries were pooled at equimolar concentrations and sequenced on an Illumina NovaSeq platform (GeneWiz) to a depth of approximately 300 million reads. *scRNAseq data annotation and analysis.* We identified 9 major cell types based on the expression patterns of marker genes. Specifically, we used *SOX2, PAX6, NES, SLC1A3* as radial glia (RG) markers, MKI67 in RG as a proliferative marker of activated RG (aRG), *ASCL1* as intermediate progenitor cells (IPC) marker, *DCX* as immature neuron marker, *GAD1* and *GAD2* as inhibitory neurons (IN) marker, *SLC17A6*, *LHX9* as excitatory neurons (EN) markers, *RELN* as Cajal Retzius neuron (CJReN) marker, and *LHX5, RELN*, as CjReN-immature neuron markers. The annotated Anndata were converted to a Seurat object. The DEG analysis was performed using the Seurat function FindMarkers (min.pct = 0.1) with the MAST method (v1.24.1). With the output from FindMarkers, significant DEGs were identified if the absolute log2 fold change was >0.58 and an adjusted p < 0.05. The pathway enrichment scores of single cells were calculated using the function ssgseaParam (minSize = 1, maxSize = Inf, alpha = 0.25, normalize = TRUE) from the R package GSVA (v1.51.14). The pseudotime analysis was performed using the R package monocle (v2.28.0). Genes related to neural development in organoids were manually selected for calculating the pseudotime.

### ScRNAseq data preprocessing

We generated gene-level UMI counts for each sample using the ParseBioscience pipeline (V1.1.2), with the GRCh38 human genome and Ensembl v108 gene annotation. We ran the remove-background module from CellBender (0.3.0) to perform ambient RNA correction. quickPerCellQC from the scuttle R package (v1.12.0) was used to identify low-quality cells based on frequently used QC metrics, including the number of genes, UMI counts, and mitochondrial DNA (mtDNA) UMI counts. Each sample was manually reviewed and we set sample-specific cutoffs to further identify high quality cells for downstream analysis. This was an essential step for single cell RNA-seq data generated from organoids where both cell sizes and cell states may differ between batches or studies. The high-quality count matrices were then converted into the Anndata format to be further analyzed using the Scanpy workflow.

Specifically, raw counts were normalized and log transformed using the normalize_total and log1p functions. We computed the top 2000 highly variable genes using the highly_variable_genes function. The highly variable features were used to perform principal component analysis. Integration was performed with Harmony using the first 25 principal components, from which we generated a neighborhood graph and dimensionally reduced representation of our data through UMAP. Clusters were identified using the unsupervised graph-based clustering algorithm, Leiden (v0.10.2) (repository: GSE304672, processed and annotated data DOI: 10.17632/58dgjxfwyv.1).

Differentially expressed genes identified from single-cell transcriptomic datasets were analyzed to uncover functionally enriched pathways across distinct cell populations. To investigate metabolic alterations, gene symbols were mapped to the Kyoto Encyclopedia of Genes and Genomes (KEGG) database to identify significantly overrepresented metabolic modules. In parallel, genes associated with DNA damage and repair were curated from published studies^113–115^ and the Gene Set Enrichment Analysis (GSEA) Molecular Signatures Database (MSigDB) (GOBP_DNA_REPAIR; https://www.gsea-msigdb.org/gsea/msigdb/cards/GOBP_DNA_REPAIR.html, https://www.gsea-msigdb.org/gsea/msigdb/cards/KEGG_HOMOLOGOUS_RECOMBINATION, cell senescence GO term: https://www.gsea-msigdb.org/gsea/msigdb/human/geneset/GOBP_CELLULAR_SENESCENCE.html) for targeted evaluation of DNA repair–related processes.

### RNA extraction and qRT-PCR

RNA from cell cultures was harvested using either the Qiagen miRNeasy (Qiagen #217004) or Qiagen RNeasy (Qiagen #74104) according to the manufacturer’s protocol, including on-column DNase digestion according to the manufacturer’s protocol (Qiagen #79254). From the purified RNA, 1–3 μg total RNA was used to perform reverse transcription cDNA synthesis using the M-MLV reverse transcriptase kit with random hexamer primer (Invitrogen #2802013) according to the manufacturer’s protocol. qRTPCR was performed using a CFX96 Real-Time System (Bio-Rad) using PowerUp SYBR Green Master Mix (ThermoFisher #A25741), 0.4 μM forward and reverse primers, and 1:20 dilution of cDNA. The following cycling conditions were used: 95◦C for 5 min, 39 cycles of 95◦C for 11 s, 60◦C for 45 s, plate read, a final melt of 95◦C, and melt curve of 65–95◦C at +0.5◦C increments. The specificity of the amplification products was verified using melt-curve analysis. The Ct values were calculated with the Bio-Rad CFX Maestro Software, and relative gene expression was calculated using the DDCt method using one of the following housekeeping genes for normalization: GAPDH, UBE2D2 (**Extended Data Table 23**). All reactions were performed in technical triplicate with a minimum of three biological replicates. Data are presented as mean ± sem in figures.

### Proteomics

#### Cortical organoids

Brain organoids samples were sent to Mass Spectrometry Proteomics Core at BCM for proteomics profiling (MDS n=2, MDS treated with ASO n=3, Controls=3, Cobtrols treated with ASO=2). Brain organoids tissues were washed two times with PBS and snap-frozen. Cell pellets were lysed by 1 % SDS lysis buffer and further prepared using a modified version of the Single-Pot Solid-Phase-enhanced Sample Preparation (SP3) protocol ^116^. The sample preparation for proteome profiling including digestion, peptide desalting, and offline fractionation was similar as described before ^117^ (**Extended Data Table 24**).

#### Human CSF proteomics sample preparation

The TMT proteomics analysis was performed by Proteome Sciences plc using TMTcalibrator™ workflow (https://www.proteomics.com/services/tmtcalibrator-workflow) which was developed by Proteome Sciences to quantify low-abundance peptides in matrices with a high degree of biological complexity based on a tissue trigger/calibrant. 200 μL of each of the human CSF samples (controls n=5, MDS n=14) were depleted using the Pierce Albumin/IgG removal Kit (Thermofisher: 89875) based on the manufacturer’s protocol using 35 μl of slurry material. 13 μg of each depleted CSF sample and 2 human brain lysate (*Calibrator/Trigger brain sample*) was reduced, alkylated, and digested with trypsin to generate peptides. After desalting on SepPak tC18 cartridges, samples were lyophilized to dryness. Dry peptides from depleted CSF and brain samples were dissolved in labelling buffer. Peptides were mixed with their respective TMTpro™ 18plex reagent. Two TMTpro™ 18plex samples containing 14 depleted CSF analytical samples and 4 pooled brain digests were generated in a protein mass ratio of 1:1.75. 300 μg from each of the two TMTpro™ 18plex samples were fractionated to finally generate 15 fractions. The fractions were lyophilised to dryness and stored at −80°C until LC-MS/MS analysis (**Extended Data Table 25**).

#### Liquid Chromatography Mass Spectrometry (LC-MS/MS)

Each of the individual fractions was analyzed by LC-MS/MS using an EASYnLC-1000 system coupled to an Orbitrap FusionTM TribridTM Mass Spectrometer (both Thermo Scientific). Samples were loaded to trap column and resolved using an increasing gradient of 0.1%FA in 80% ACN through a 50 cm 75 um ID EasySpray analytical column (Thermo Scientific, PN ES803A) at a flow rate of 300 nL/min. The mass spectrometer was operated in data dependent mode with full scans at 120,000 resolution and MS2 fragment scans acquired at 50,000 resolution with the automatic gain control (AGC) setting set to A-C4. All mass spectrometry data files were processed in Proteome Discoverer (PD) v2.5 (Thermo Scientific). The enzyme specificity was set to trypsin, with the search allowing for up to 2 missed cleavages. The precursor mass tolerance was set to 10 ppm, while the fragment tolerance was set to 0.02 Da. TMTproTM modification of N-termini and lysines, and carbamidomethylation of cysteines were set as static modifications, while methionine oxidation was set as a variable modification.

### CUT & RUN data analysis

We downloaded a previously published CUT&RUN data (GEO accession id GSE213752) performed for *mecp2* in *mecp2*-knockout, wild-type, and Mecp2-transgenic hippocampus of Mus musculus, accompanied with IgG controls (**Fig. 3D** and **Extended Data Fig 6E-G**).^118^

The data was analyzed using the nf-core cutandrun workflow (v3.2.2). Reads were mapped to the mouse genome (mm10) and the spike-in genome (K12). Peaks were identified using the SEACR ^119^algorithm. We further identified differential peaks by comparing the WT samples to the *mecp2*-knockout samples using DiffBind^120^(v3.12). We used wiggletools^121^(v1.2.11) to generate averaged signals for each genotype. Summary plots were generated using R package Gviz (v1.4.61).

### Immunostaining

Human brain organoids were fixed using 4% paraformaldehyde (PFA) in PBS for 3 hours. min. After fixation, organoids and primary tissue samples were washed in PBS (pH7.4), cryoprotected in 30% sucrose, washed with PBS, embedded in Optimal Cutting Temperature sectioning medium, and cryosectioned at 14 µm of section thickness on glass slides. For MSI organoids sections were embedded, fixed and stained as previously described^36^.

For disaggregated organoids, cells were plated onto PhenoPlate 96-well microplates 96wells (50-209-9831) or 4-well Millicell EZ SLIDE glass, sterile, EMD Millipore PEZGS0416, 16/PK, coated with Poly-D-lysine hydrobromide,mol wt >300,000 (50 ug/ml) and mouse laminin (25 ug/ml), recover for 4 dfays and then fixed with PFA for 20 mins following by washes in PBS. Subsequently, sections/cells were permeabilized using 0.1% Triton X-100 for 20 minutes, followed by blocking in a solution containing 5% donkey serum (Stratech, 017-000-121-JIR), 2% (w/v) BSA, and 0.1% Triton X-100 in PBS for 1 hour. For primary antibody staining, the following antibodies in blocking solution were used at 4°C overnight: anti-beta III tubulin (TUBB3) (Sigma-Aldrich, #mab1637, 1:200), anti-TUBB3 (Biolegend, #801202, 1:500), anti-MAP2 (Fisher Scientific, MAB3418MI, 1:200), anti-GFAP (Thermo, #13-0300, 1:100), anti-S100B (DAKO, #ZO311; 1:100), anti-SOX2 (R&D system, #AF2018, 1:200), anti-PAX6 (ThermoFisher, #42-6600, 1:100), anti-DCX (Santa Cruz, #SC-271390, 1:150), anti-Ki-67 (Cell Signaling, #9129S; 1:300), anti-TOM20 (Santa Cruz Biotechnology, #sc-17764, 1:200), anti-UQRCQ (Proteintech, #14975-1-AP, 1:200), anti-PFAS/FGAM (Bethyl Laboratories, #A304-218A, 1:250), anti-GART (Novus Biologicals, # H00002618-M01, 1:200), anti-Phosphorylated Vimentin (MBL Life science, #D076-3, 1:500), anti-MECP2 (Cell Signalling, #3456, 1:80), anti-SDHB (Abcam, #21A11AE7, 1:250), anti-8-hydroxyguanosine (Millipore Sigma, #DR1001, 1:100), Alexa Fluor® 647 anti-H2A.X Phospho (Ser139) (Biolegened, #613407, 1:100), Alexa Fluor® 647 anti-COXIV (Cell Signalling, #7561, 1:200). After washing off the primary antibodies, the following secondary antibodies in washing buffer were used for 2 hours: donkey anti-goat Rhodamine Red™-X (RRX) (Jackson ImmunoResearch, #705-295-003, 1:500), donkey anti-rabbit Alexa 488 (Abcam, # ab150073, 1:500), donkey anti-rabbit Alexa 594 (Abcam, # ab150076, 1:500), donkey anti-rabbit Alexa 647 (Jackson ImmunoResearch, # 711-605-152, 1:500), donkey anti-rat Alexa 647 (Jackson ImmunoResearch, # 712-605-153, 1:500), donkey anti-rat Alexa 488 (Abcam, # ab150153, 1:500), donkey anti-mouse Alexa 488 (Abcam, # ab150105, 1:500), donkey anti-mouse RRX (Jackson ImmunoResearch, #715-295-151, 1:500), donkey anti-mouse Alexa 647 (Jackson ImmunoResearch, #715-605-150, 1:500).

For additional DNA staining sections/cells were incubated with DAPI (1 µg/ml) for 10 min at room temperature. The DAPI solution was washed off twice with PBS before mounting the coverslips on glass slides with the anti-fading reagent (Prolong Diamond Antifade mounting medium, Thermo Fisher Scientific). For super-resolution STED microscopy, the blocking buffer was prepared using 5% goat antiserum to match the host species of the secondary antibodies raised in goat. Indeed, secondary STAR ORANGE goat anti-mouse (Fisher Scientific, NC1933863, 1:200), STAR RED goat anti-rabbit, (Fisher Scientific, NC1933870, 1:200) were used for STED. For MSI, Histological sections extracted from each block underwent routine Hematoxylin eosin staining staining following standard protocol ^122^. Commercially available Instant Hematoxylin from Thermo scientific® was used.

To assess the levels of neurogenesis in brain organoids we labeled the dividing neural progenitor cells by 5-ethynyl-2’-deoxyuridine (EdU) and analyzed their neuronal differentiation 7 days after EdU labeling. Proliferating cells were labeled using the Click-iT™ EdU Alexa Fluor™ 488 Imaging Kit (Fisher Scientific, #C10337) according to the manufacturer’s protocol and they were co-stained with DCX a marker of newborn neurons doublecortin to check the number of newly differentiated neurons (Edu/DCX+). Confocal z-stacks from tissue sections were acquired using a Leica SP8 microscope with 10× and 63× objectives. Cells cultured on plates were imaged at 60× and 100× magnification using a Nikon spinning disk microscope. Single-plane images from maximum intensity projections are shown. Live imaging was performed using the Yokogawa CV8000 high-content imaging system. Super-resolution microscopy was performed using a 3D Stimulated Emission Depletion (STED) Super Resolution System (Abberior 3D STED), achieving a resolution of approximately 50 nm, sufficient to visualize subcellular structures such as purinosomes. All images were processed using Fiji (version 2.1.4) and TRUESHARP for STED-specific image enhancement. Mitochondrial morphology and network organization were analyzed using the Mitochondria Analyzer plugin in combination with a deconvolution tool for semi-automated, image-based quantification in both 2D and 3D. Morphological and network parameters were computed at both the per-cell and per-mitochondrion level in control and MDS samples. Neurite Tracer and Skeleton (2D/3D) plugins in Fiji were used for neuromorphometric analysis. Individual branches were traced, and quantitative analyses were performed on a per-neuron basis.

### Western Blot

Protein expression was quantified using the Wes™ Simple Western system (ProteinSimple, Bio-Techne) according to the manufacturer’s instructions. Human brain organoids were first washed with ice-cold PBS, then lysed in ice-cold RIPA buffer (50 mM Tris-HCl pH 7.5, 150 mM NaCl, 1% Triton X-100, 0.5% sodium deoxycholate, 0.1% SDS, 5 mM EDTA) supplemented with phosphatase inhibitor cocktail (GenDEPOT #P32000), protease inhibitor cocktail (GenDEPOT #P3100), and Pierce™ Universal Nuclease for Cell Lysis (ThermoFisher #88700; 1:1000 dilution). Lysates were rotated for 20 minutes at 4°C, centrifuged at maximum speed for 20 minutes at 4°C, and the supernatant was collected. Protein concentration was determined using the Pierce™ BCA Protein Assay Kit (ThermoFisher #23225). For Wes analysis, protein samples were mixed with Simple Western Sample Buffer and ProteinSimple MasterMix (containing DTT and fluorescent standards), and loaded into 25-capillary assay plates along with primary antibodies, HRP-conjugated secondary antibodies, separation and stacking gel matrices, and luminol-peroxide substrate. Separation and chemiluminescent detection were performed automatically, and signal was quantified using Compass software as electropherogram peak area and visualized in gel view. Antibodies used were PFAS (Bethyl Laboratories, #A304-220A, 1:80), β-Actin (Cell Signaling, #4967S, 1:10), and MeCP2 (Cell Signaling, #3456S, 1:50).

### Functional Mitochondrial validation

#### Real time Glutathione assay

Basal mitochondrial glutathione (GSH) levels were first measured in real time using the Mito-RealThiol (MitoRT) probe (EBY003) at 1 µM, following the protocol described previously^123^. In a separate set of experiments, cells were exposed to oxidative stress by treatment with 100 µM hydrogen peroxide (H₂O₂), and GSH levels were measured after the stressor. This ratiometric assay enabled quantification of reduced GSH in both basal and stressed conditions across experimental groups. Live-cell imaging was performed using the Yokogawa CV8000 high-content imaging system, and image analysis was conducted with ImageJ software.

#### Mitochondrial dyes

Using DMSO, the MitoTracker probes were diluted to 1mM. Cells were incubated 200 nM of either MitoTracker Red CMXRos (Ex 579nm; Em 599nm) or MitoTracker Green FM (Ex 490 nm; Em 516 nm) for 30 min at 37C. Following incubation, cells were washed 3 times in pre-warmed, equilibrated medium and either fixed in 4% PFA (MitoTracker Red CMXRos) for further staining or for flow-cytometry (MitoTracker Green FM).

Tetramethylrhodamine, Ethyl Ester, Perchlorate (TMRE) (Invitrogen, Cat# T669) (Ex 559 nm; Em 574 nm) and CellRox Deep Red (Em 644, Ex 665, Fisher scientific, Cat#C10422) (Ex 559 nm; Em 574 nm) or CellROX green (Em 485, Ex 520 nm, Fisher scientific, Cat#C10444) were used at a concentration of 400 nM and 5 uM respectively to stain cells incubated for 30 minutes at 37°C for live-cell imaging and/or flow cytometry.

#### Mitochondrial drugs

Coenzyme Q10 (ubidecarenone; Thermo Scientific, #J65137-03) was used at a final concentration of 5 μM for 7 days, with media changed every other day, as previously described^124^. Oligomycin (Selleck Chemicals, #S1478) was applied at 1 nM in organoid media for one week following established protocol^125^. Additionally, 2S-octylhydroxyglutarate (Fisher Scientific, #501968150) was used at 100 μM in control organoids for 10 days, as previously described^35126^.

### Oxygen consumption rate measurement (OCR)

#### Seahorse

The Seahorse XF96 Mito Stress Test was used to measure oxygen consumption rate (OCR) in disaggregated organoids seeded at a density of 80,000 cells per well on poly-D-lysine and laminin-coated XF96 plates. After 72 hours of plating, metabolic measurements were performed using the Seahorse XF96 Extracellular Flux Analyzer. Prior to the assay, the culture medium was replaced with Seahorse XF DMEM medium (pH 7.4) supplemented with 21 mM glucose, 2 mM glutamine, and 1 mM pyruvate. OCR was measured under basal conditions, followed by sequential injections of 1 μM oligomycin, 1 μM FCCP, and a combination of 0.5 μM antimycin A and rotenone. Data were normalized to cell number using nuclear counterstaining. *Resipher.* Resipher employs an optical measurement-based method to assess Oxygen Consumption Rate (OCR) in cell cultures, enabling long-term, non-invasive, real-time monitoring of oxygen consumption directly within standard cell culture plates and incubators^127^. Cells from disaggregated brain organoids were plated at a seeding density of 50,000 cells per well, and basal OCR was continuously measured for 3 days in control and MDS-derived organoid cells. Additionally, OCR was assessed following specific inhibition of Complex III using antimycin A (0.1 μM)^128^, and upon treatment with Coenzyme Q10, as previously described.

### Flow-cytometry

Cells derived from disaggregated organoids were washed twice with cold PBS, filtered to obtain single-cell suspensions, and stained with TMRE, MitoTracker, or CellROX as indicated. Data were acquired using a BD FACSCelesta cell analyzer. Mean fluorescence intensity was measured in at least triplicate samples and analyzed with FlowJo software. For all flow cytometry analyses and sorting, DAPI staining was used to assess DNA content or exclude dead cells, depending on the experiment. Gating strategies consistently included exclusion of doublets and DAPI-positive dead cells.

### MECP2 Binding Motif Analysis

The MECP2 binding motif “SCCGGAG” was retrieved from the AnimalTFDB database (accessed September 2025) and corresponds to the consensus MECP2 motif annotated in HOCOMOCO v11 (MECP2_HUMAN.H11MO.0.C). The human genome sequence (GRCh38/hg38) was obtained using BSgenome.Hsapiens.UCSC.hg38 (v1.4.5), and the 1,000 bp upstream region of each annotated transcription start site (TSS) was extracted to define promoter sequences. Motif scanning was performed using FIMO (MEME Suite v5.5.5) with default parameters, and genes were classified as putative MECP2 targets if at least one occurrence of the canonical motif or its variants (e.g., “GCCGGAG,” “CCCGGAG”) was detected, yielding 7,323 candidate targets; multiple motif occurrences within a single gene were recorded but not counted separately. To explore the metabolic context of MECP2 binding, we analyzed the top 10 KEGG pathways enriched in the metabolomic dataset, including citrate cycle (TCA cycle, hsa00020), oxidative phosphorylation (hsa00190), starch and sucrose metabolism (hsa00500), nicotinate and nicotinamide metabolism (hsa00760), propanoate metabolism (hsa00640), nitrogen metabolism (hsa00910), glutathione metabolism (hsa00480), purine metabolism (hsa00230), glycolysis/gluconeogenesis (hsa00010), and glycerophospholipid metabolism (hsa00564). These pathways collectively comprised 726 unique genes, of which 178 overlapped with the MECP2 motif gene set; among these, 38 belonged to the TCA cycle and oxidative phosphorylation pathways, 35 to glycerophospholipid metabolism, 32 to purine metabolism, and 17 to glutathione metabolism (**Extended data Table 26)**.

### Mitomics data analysis

Processed data from the Mitomics Project was downloaded from https://www.mitomics.app/DV/J51Lhy2/main.php. Both two metabolomics datasets (Metabolomics_3Reps and Metabolomics_Batch3and4) were used to visualize the z-scores of tricarboxylic acid (TCA) cycle and purine metabolites.

Metabolites were identified and clustered based on annotations from the KEGG and HMDB databases. Correlation and regression analyses were performed to assess relationships between metabolite levels and experimental variables.

## Extended Data TABLES

**Extended Data Table 1** Clinical data from patients

**Extended Data Table 2** MDS patients genomic coordinates

**Extended Data Table 3** Drugs from patients

**Extended Data Table 4** Extended Data Fig. 2 A-B metabolites from heatmap

**Extended Data Table 5** Human blood dataset metabolome

**Extended Data Table 6** Human blood dataset metabolome from GMAP

**Extended Data Table 7** Mice blood dataset metabolome including aso

**Extended Data Table 8** Combo metanalysis blood dataset

**Extended Data Table 9** Combo metanalysis blood +aso

**Extended Data Table 10** Details of Individuals’ iPSC included in this study

**Extended Data Table 11** Neuro dataset CSF metabolome

**Extended Data Table 12** Neuro dataset brain organoids metabolome

**Extended Data Table 13** Neuro dataset mice cortex including aso

**Extended Data Table 14** Combo metanalysis neuro dataset

**Extended Data Table 15** Combo neuro dataset + aso

**Extended Data Table 16** Integrative Pathways analysis for CSF

**Extended Data Table 17** Integrative Pathways analysis for brain organoids

**Extended Data Table 18** Integrative Pathway + enrichment analysis CSF – Byocic

**Extended Data Table 19** Integrative Pathway + enrichment analysis organoids – Biocyc

**Extended Data Table 20** TCA metabolites analysis mitomics

**Extended Data Table 21** Hydroxyglutaric acid (Molecule)_LookupData

**Extended Data Table 22** MSI parameters used

**Extended Data Table 23** Primers used for qPCR

**Extended Data Table 24** Proteome organoids

**Extended Data Table 25** Proteome CSF

**Extended Data Table 26** MECP2 prediction binding site

**Extended Data Video 1 | Three-dimensional reconstruction of mitochondrial networks in control and MDS astrocytes.** 3D rendering of mitochondrial volumes reveals altered morphology in MDS astrocytes, characterized by reduced branching, fragmentation, and smaller organelle size compared to controls.

