## Supplemental Figures for "MECP2 Duplication Uncouples Mitochondrial and Purine Metabolism During neuronal maturation"

Extended Data Fig. 1

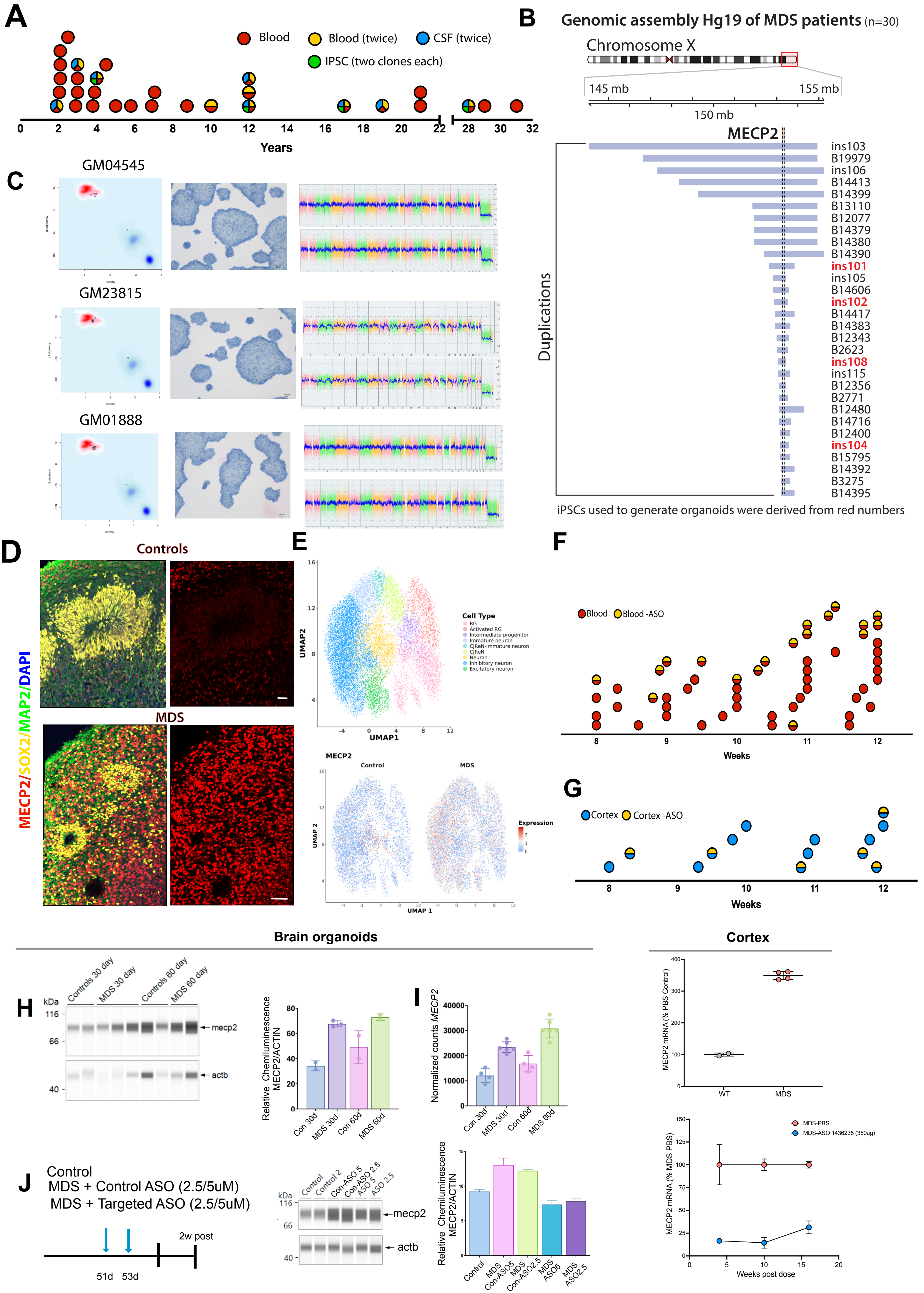

Extended Data Fig. 2

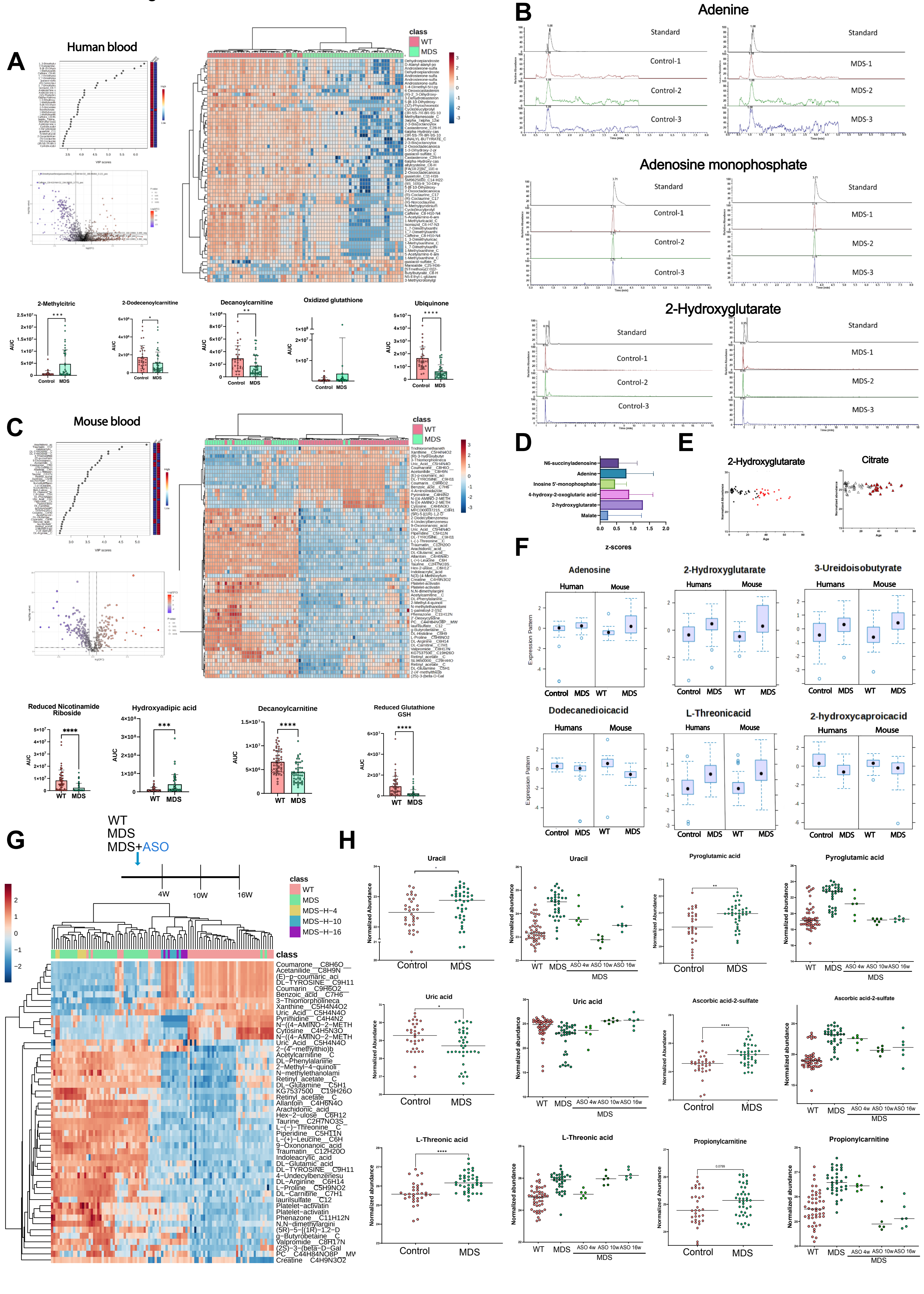

Extended Data Fig. 3

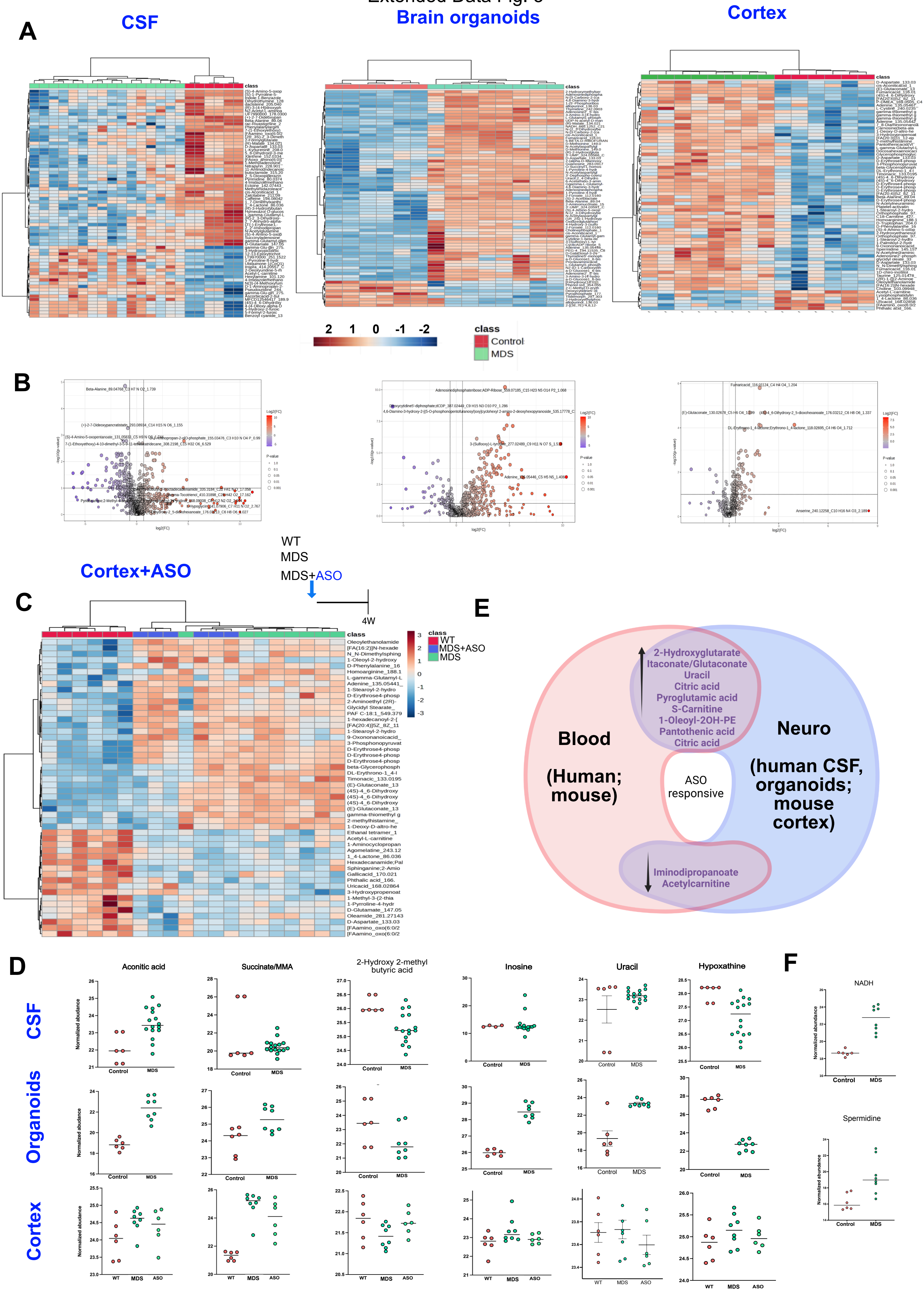

### Extended Data Fig.4

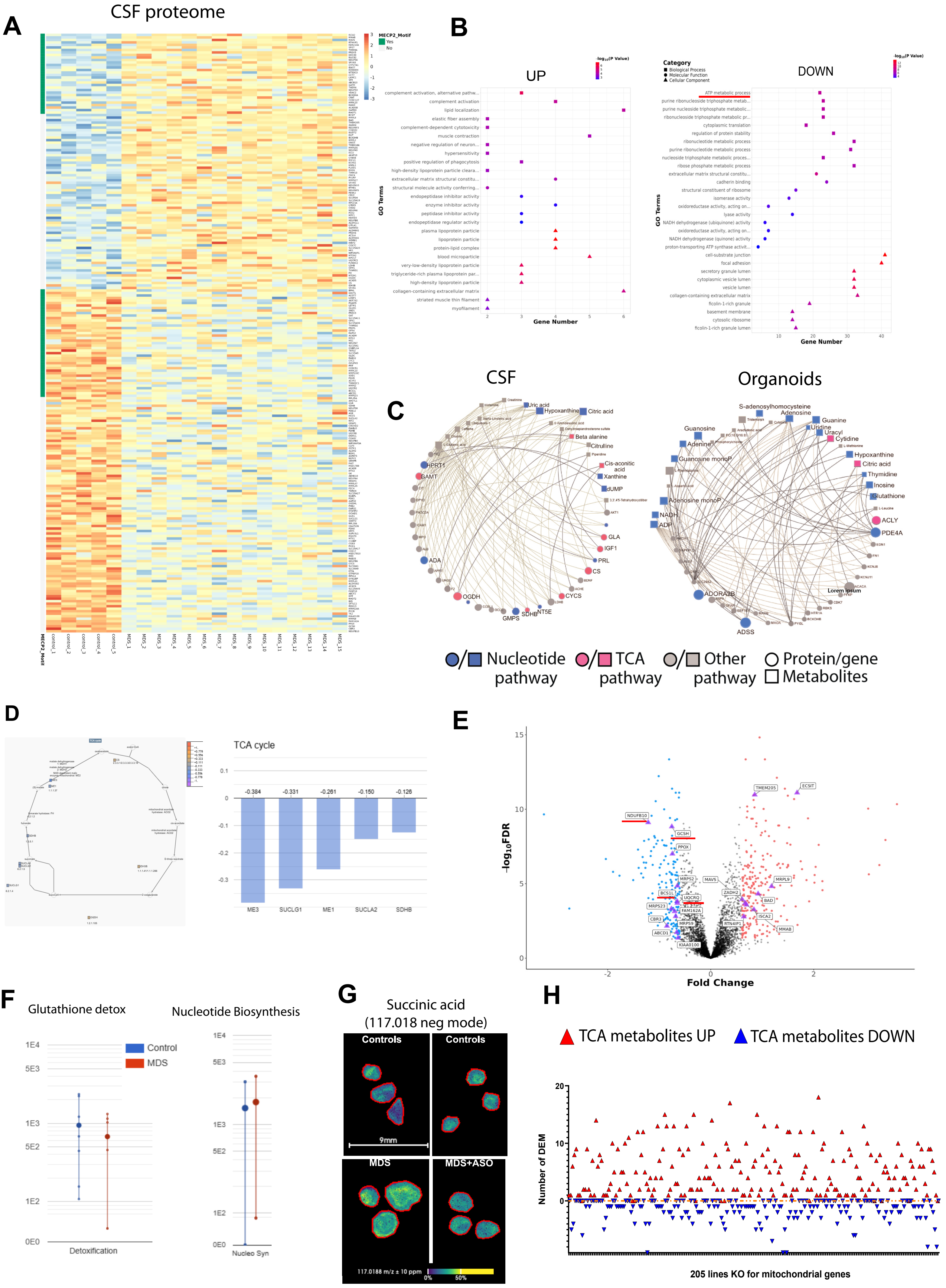

Extended Data Fig. 5

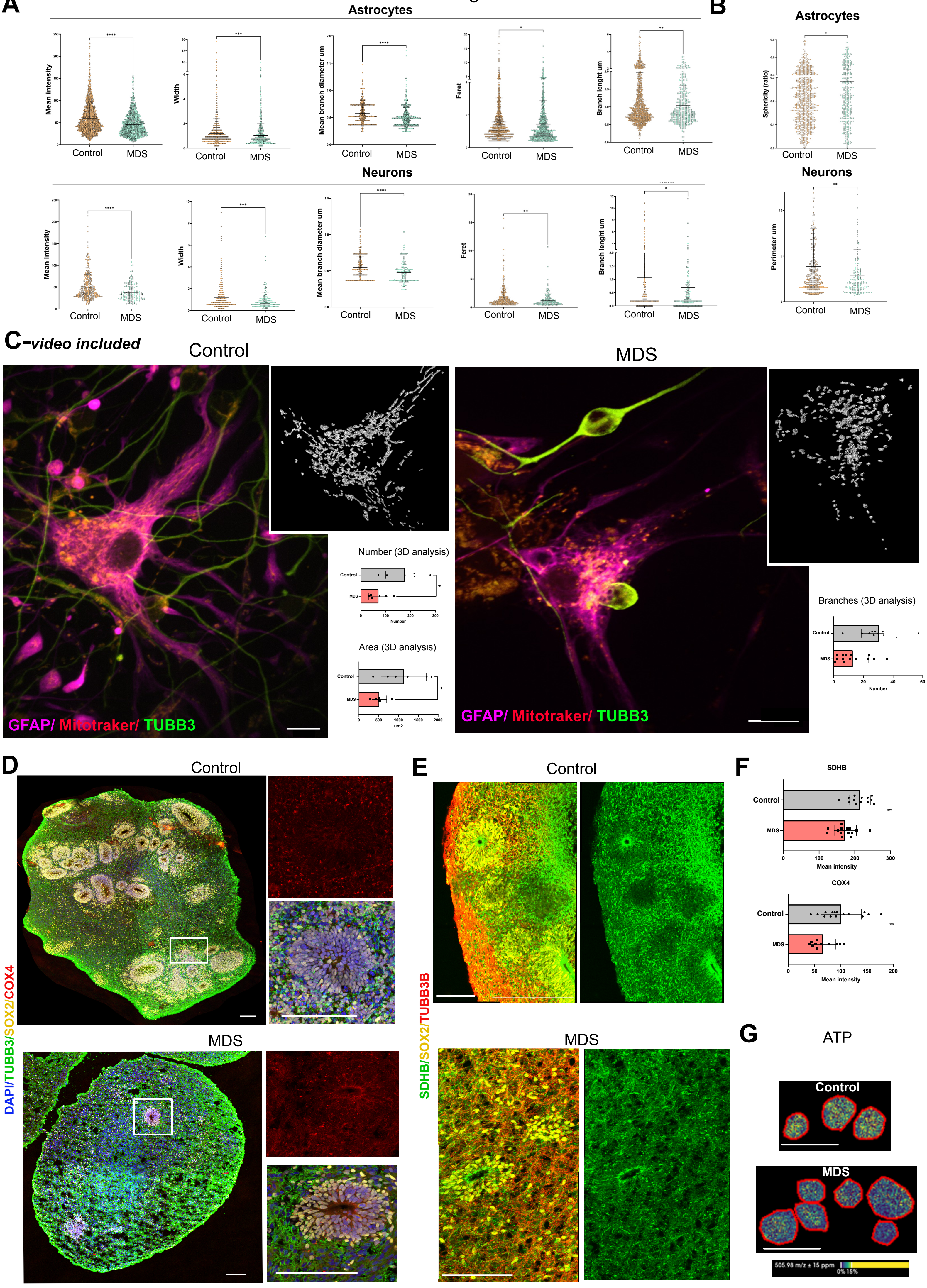

### Extended Data Fig. 6

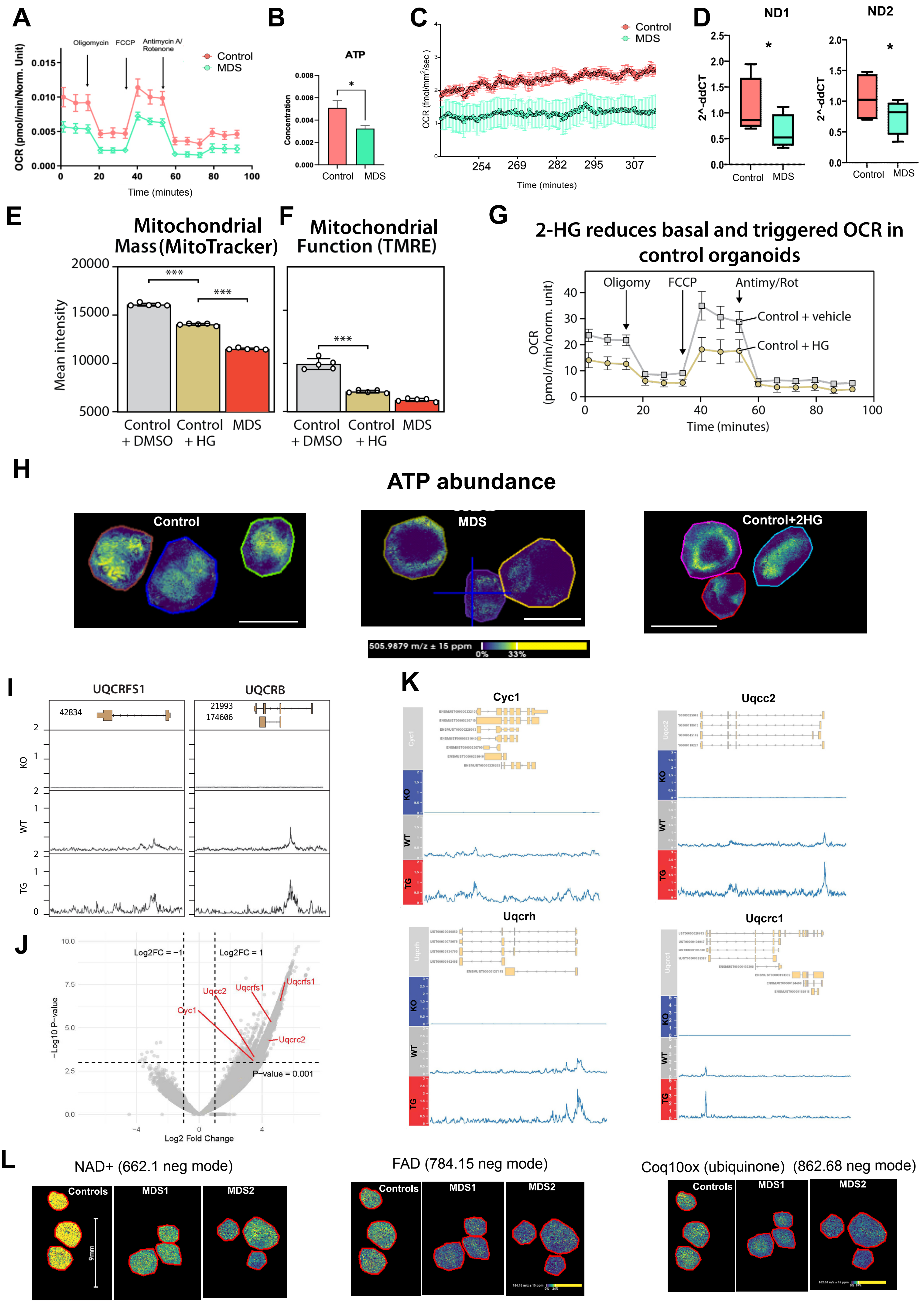

Extended Data Fig. 7

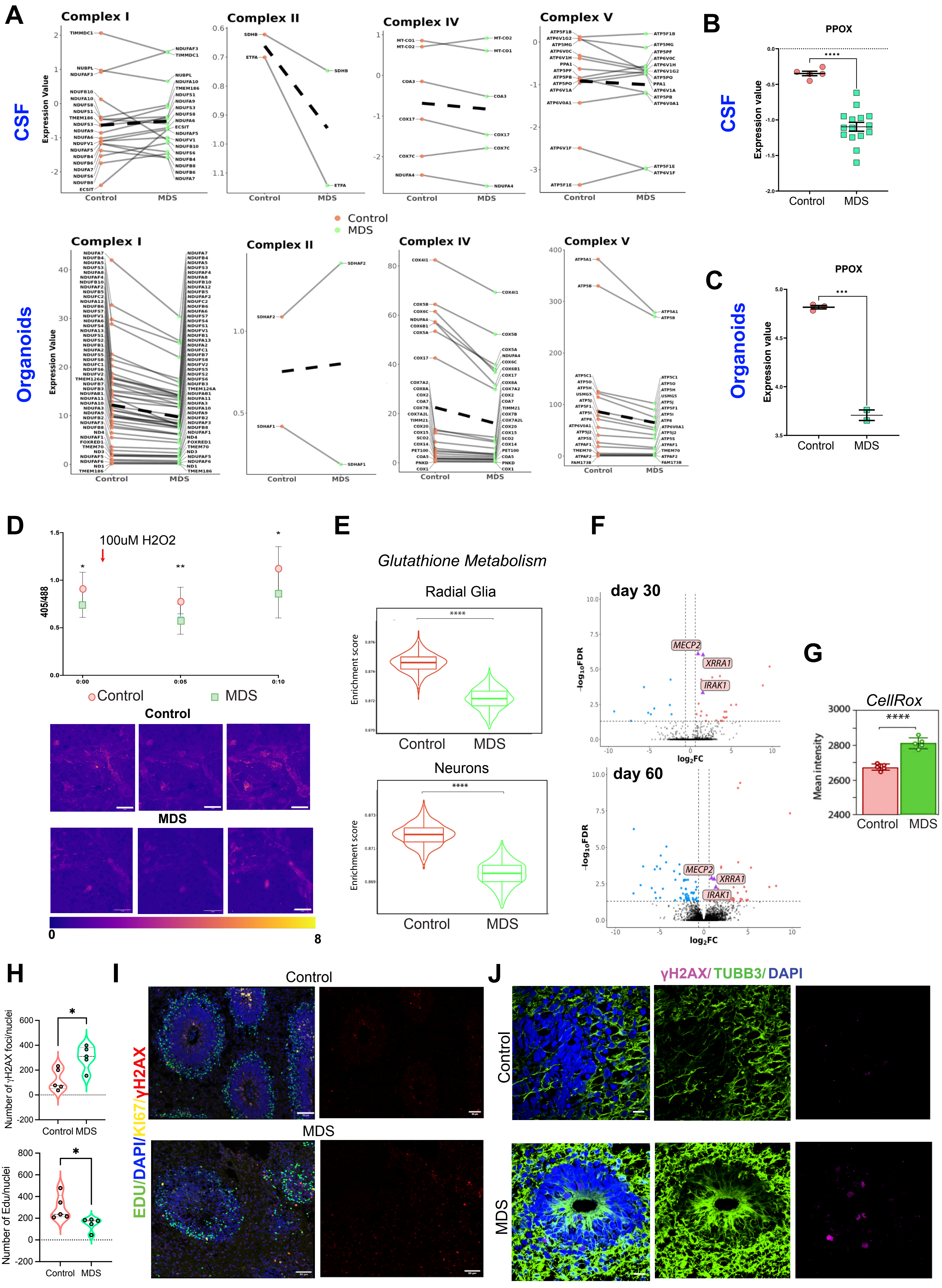

Extended data Fig. 8

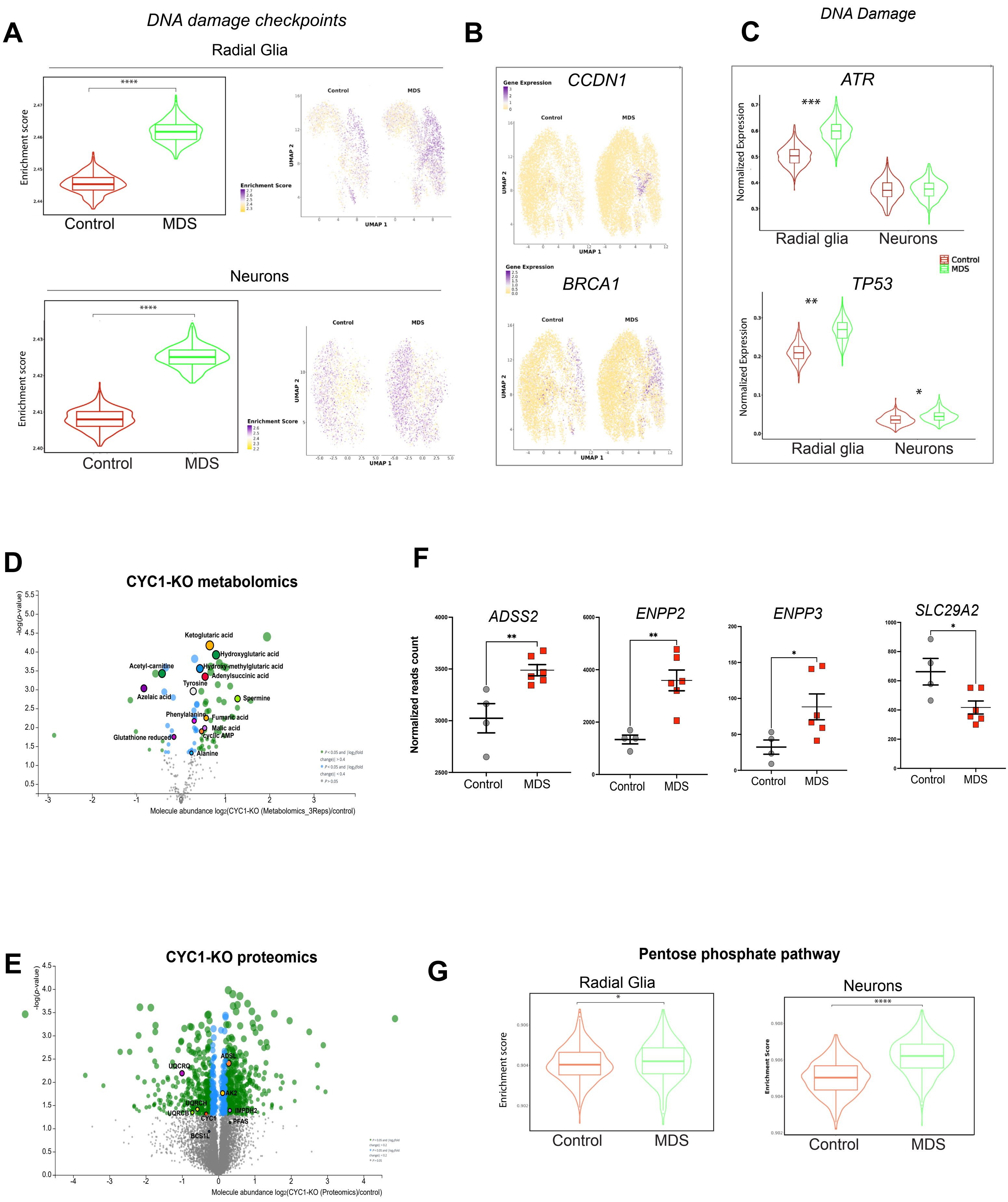

### Extended Data Fig. 9

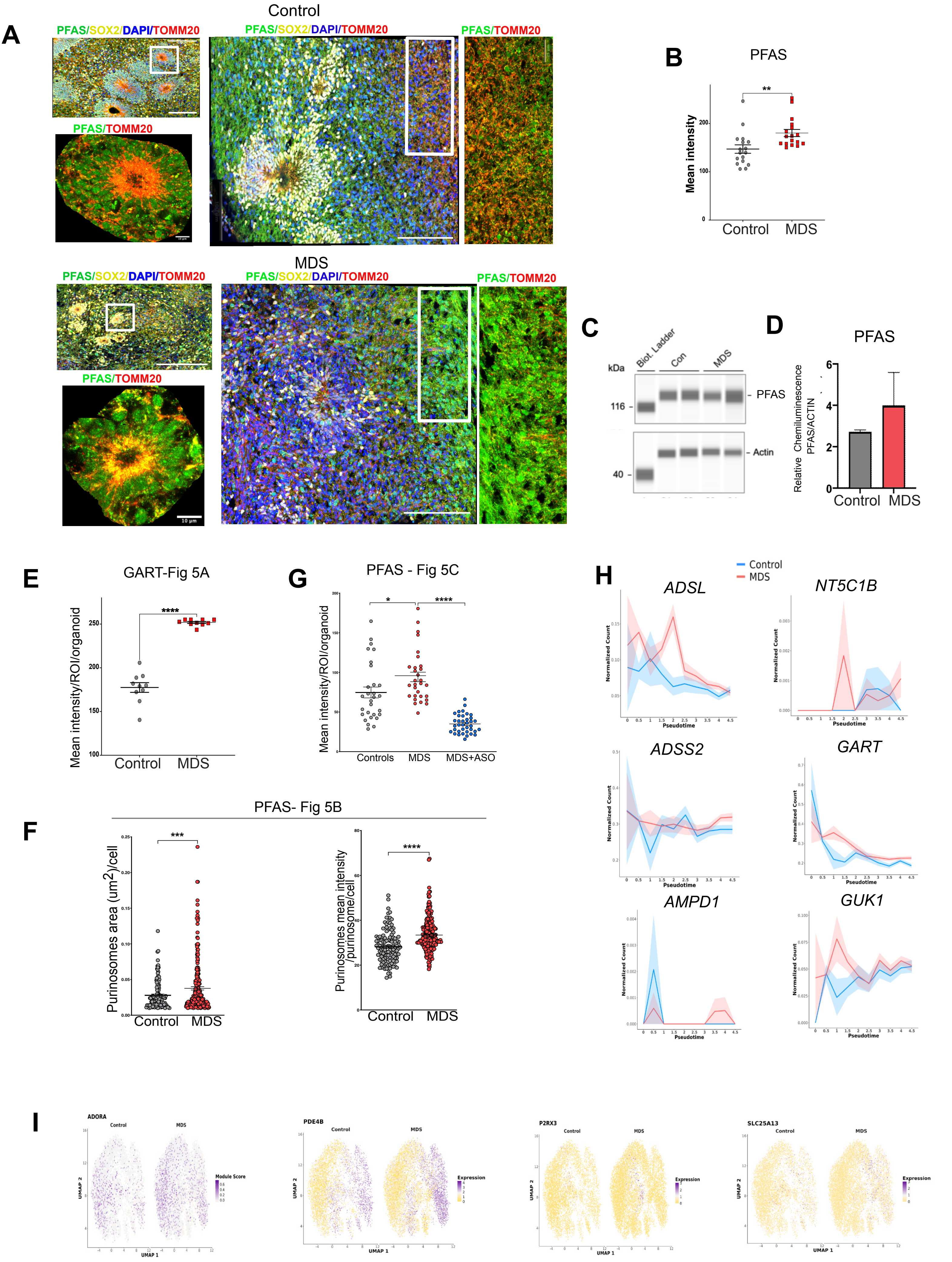

Extended Data Fig. 10

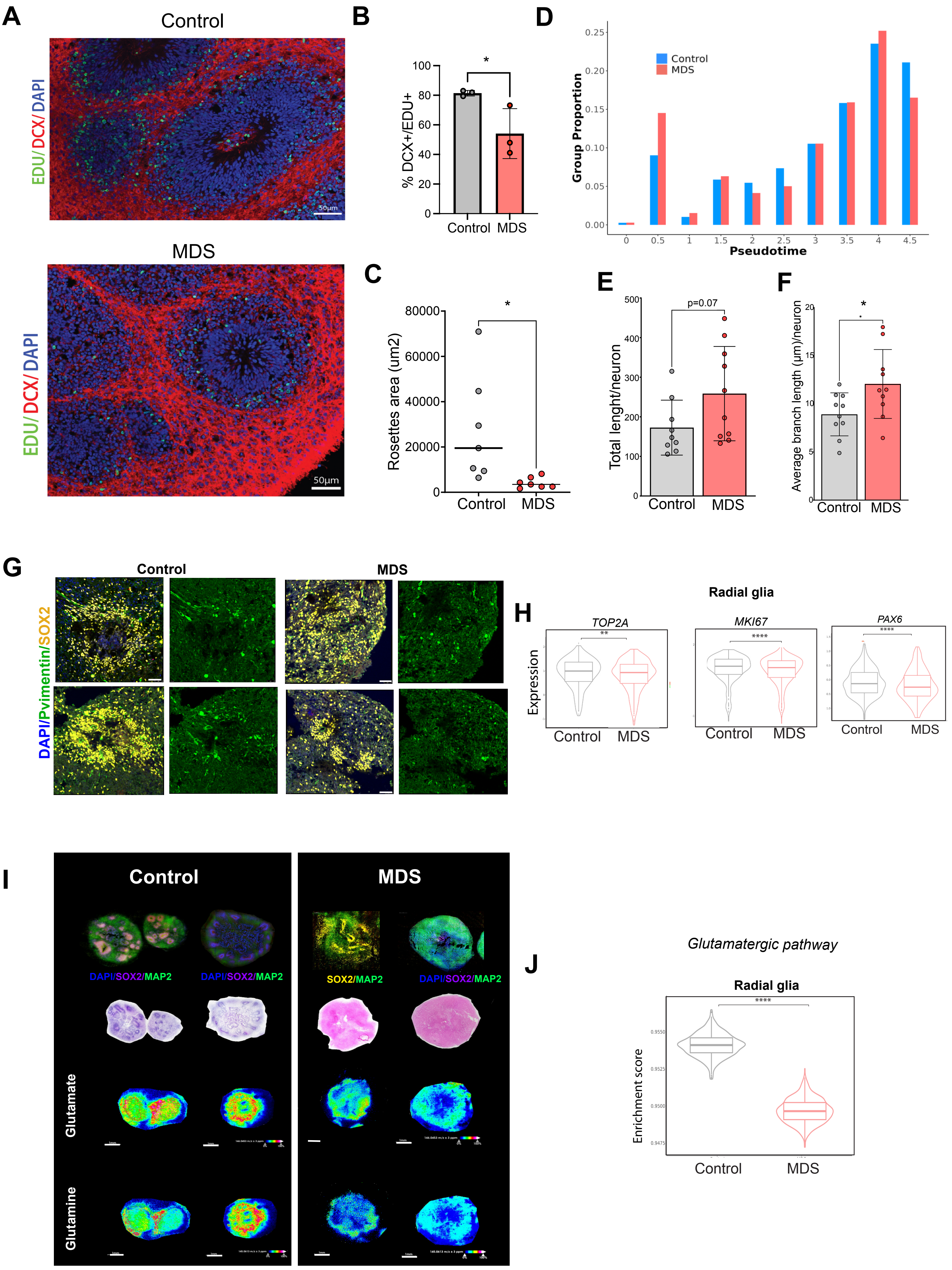
